# Astrocytic Gapjinc (TMEM43) modulates gap junction networks by facilitating transjunctional potentials

**DOI:** 10.1101/2022.11.08.515259

**Authors:** Minwoo Wendy Jang, Ki Jung Kim, Jiwoon Lim, Daeun Kim, Yongmin Mason Park, Tai Young Kim, Woo Suk Roh, Young-Eun Han, Joungha Won, Jae-Hun Lee, Eunjin Shin, Ah-reum Han, Ho Min Kim, Kyung-Seok Han, C. Justin Lee

## Abstract

The *TMEM43* gene has been reported to play supportive but critical roles in human diseases including cancer, arrhythmogenic right ventricular cardiomyopathy (ARVC), and auditory neuropathy spectrum disorder (ANSD). However, direct characterization of the TMEM43 protein itself and its role in the brain remain unexplored. In this study, we demonstrated that TMEM43 confers ion channel activities via the lipid bilayer reconstitution of purified TMEM43 protein and further characterized TMEM43 as a pH-sensing cation channel in the heterologous expression system. TMEM43 was shown to conduct transjunctional potentials between adjacent cells, further facilitating electrical couplings of the gap junctions. In the hippocampus of TMEM43 knockout (KO) mice, we observed a decrease in astrocytic dye diffusion and potassium buffering, an increase in neuronal excitability, and alterations in AMPA/NMDA ratio and LTP. The electrophysiological changes in the KO mice led to a disturbance in memory retrieval which was rescued with TMEM43 overexpression. These results indicate that TMEM43 actively participates in gap junction networks of the hippocampus to prevent neurons from hyperexcitability, which is critical for memory retrieval. Together, our study elucidates the molecular and functional identities of TMEM43 and underscores its role in memory retrieval.

## Introduction

The *TMEM43* gene has been implicated in many human diseases such as glioblastoma^1^, pancreatic cancer^2^, arrhythmogenic right ventricular cardiomyopathy (ARVC)^3,4^, and auditory neuropathy spectrum disorder (ANSD)^5^. Mechanistically, TMEM43 interacts with the scaffold protein CARMA3 and its associating complex to induce downstream NF-κB activation and plays a critical role in controlling brain tumor progression^1^. TMEM43 is also known to stabilize PRPF3 and regulate RAP2B/ERK axis, accelerating the progression of pancreatic cancer^2^. In the heart, the *TMEM43*-p.(Ser358Leu) variant increases the stiffness of the cell nucleus, which might lead to massive loss of cardiomyocytes^6^ while altering intercalated disc protein expression, disturbing gap junction transfer, and reducing conduction velocity in cardiac cell lines^7^. In the cochlea, TMEM43 was shown to interact with the predominant gap junction channels Connexin 26 (Cx26) and Cx30 to mediate the large potassium (K^+^) conductance current in the cochlear glia-like supporting cells^5^. This K^+^ conductance was significantly reduced when the *TMEM43*-p.(Arg372Ter) variant was introduced, leading to progressive hearing loss^5^. In addition to the clinical studies, *in vitro* and *ex vivo* studies demonstrate TMEM43 interaction with secondary proteins. For instance, TMEM43 interacts with emerin and A-and B-type lamins in undifferentiated C2C12 cells^8^ and with one of the two-pore-domain potassium (K2P) channels, KCNK3 (TASK-1) in the cochlea^9^. The involvement of TMEM43 in cardiac rhythmicity^7^, gap junction-mediated dye diffusion^7^, and K^+^ current^5,9^ implies the possible role of TMEM43 in gap junction networks of facilitating ion movements.

Gap junctions are formed by two hexameric hemichannels (connexons) encoded by connexin genes, to achieve intercellular signaling. The gap junctions enable cell-cell communications via allowing transportation of ions, second messengers, metabolites, and neurotransmitters^10^. In the brain, the diverse connexins in different cell types control many essential processes such as neurodevelopment and neuroprotection^11^. The majority of gap junctions in the brain are formed between astrocytes, making an astrocytic syncytium. This astrocytic syncytium regulates neuronal excitability and network homeostasis by actively buffering K^+^ and glutamate in the extracellular space, released from activated neurons^12^. Previous studies have shown that loss of connexins result in enhanced neuronal excitability^13^, impaired memory^14,15^, and hypertrophy in astrocyte and microglia^13,14^. On the other hand, connexin expression was shown to be increased, along with reactive astrogliosis, in Alzheimer’s disease mouse models^16–18^ and in ischemia or brain injuries^19^. The diverse working mechanism of the connexins may come from different modulatory proteins. For instance, assembly of carcinoma-astrocyte gap junctions composed of Cx43 is promoted by protocadherin 7 (PCDH7), increasing cGAMP transfer to astrocytes^20^. Neuronal Cx36 interacts with calcium/calmodulin-dependent kinase II (CaMKII) and is phosphorylated, possibly enhancing synaptic efficacy^21^. It has also been reported that K^+^-induced increase in astrocyte coupling is mediated by Ca^2+^ influx through L-type Ca^2+^ channels, independent of Cx43 expression level, indicating that increase in astrocytic junctional communication is related to an increased number of active channels within gap junction plaques^22^. Indeed, many associated proteins such as enzymes, protein phosphatases and kinases, cytoskeletal elements, and membrane receptors play essential roles in regulating the assembly, function, and life-span of the connexins^23^.

Although TMEM43 has been reported to play supportive but critical roles in many human diseases, direct characterization of TMEM43 protein itself and its role in the brain remain unexplored. In this study, we disclose TMEM43 as a new class of a pH-sensing cation channel and demonstrate that TMEM43 is engaged in facilitating astrocytic diffusion system, which is required for successful memory recall.

## Results

Given the strong association of TMEM43 with ionic movements, we examined if TMEM43 can elicit single-channel currents after TMEM43 protein purification (**Fig. 1a**). Evident voltage-dependent channel openings and closings were observed in the lipid bilayer-reconstituted TMEM43 protein (**Fig. 1b-d**). TMEM43 was silent at a holding potential of 0 mV, but the channel activity increased with growing holding potentials in both positive and negative directions (**Fig. 1c,d**). The direction of the TMEM43-induced current switched at 0 mV, indicating the reversal potential of TMEM43 at around 0 mV. The stochastic channel openings observed in TMEM43 are a typical feature of an ion channel. An ion channel must traffic to the cell plasma membrane to elicit channel currents. We have previously shown that TMEM43 protein localizes to the cell surface by biotinylation assay and immunostaining^5^. To validate TMEM43 localization and topology in this study, we tagged Myc and FLAG at the N-terminus and C-terminus of TMEM43, respectively (**Fig. 2a**), and performed immunocytochemistry with or without cell permeabilization in the heterologous expression system. As expected, both Myc and FLAG signals were detected from TMEM43-expressing cells in cell permeabilization and non-permeabilization conditions, demonstrating that N-and C-terminus are at the extracellular space (**Fig. 2b**). These results also support the membrane trafficking property of TMEM43. We next investigated the channel characteristics of TMEM43. We expressed TMEM43 in CHO-K1 cells and measured its membrane currents under a voltage clamp. Compared to naïve CHO-K1 cells, TMEM43-expressing CHO-K1 cells displayed significantly higher amplitude of both inward and outward passive membrane currents with an average reversal potential of -15.5 mV, after junction potential correction (**Fig. 2c,e,f**). Furthermore, TMEM43-induced current showed a doubly rectifying I-V curve, resembling the voltage-dependent probabilistic channel openings measured from the purified TMEM43 protein (**Fig. 1b**). The inward current mediated by TMEM43 was due to an influx of sodium ions (Na^+^) based on the observation that the substitution of Na^+^-free external solution eliminated the inward current (**Fig. 2d,e**) with a significant shift of reversal potential (**Fig. 2f**). The outward current was carried by potassium ions (K^+^), as evidenced by the observation that the outward current was absent with an elimination of intracellular K^+^ (**Fig. 2d,e**) or by cesium ions (Cs^+^) as the inward current was still present after intracellular K^+^ substitution to Cs^+^ (**Extended Data Fig. 1a-f**). These results indicate that TMEM43 is a nonselective cation channel with permeability to cations such as Na^+^, K^+^, and Cs^+^. As expected, TMEM43-mediated current was significantly and potently blocked by a well-known nonselective cation channel blocker GdCl3^24^, with a half-maximal inhibition of 0.840 μM (**Fig. 2g,h**). In addition, TMEM43-mediated current was not affected by either intracellular chloride ion (Cl^-^) (**Extended Data Fig. 1a-c**) or calcium ion (Ca^2+^) (**Extended Data Fig. 1d-f**), indicating that TMEM43 is neither an anion channel nor a Ca^2+^-activated channel. It is noteworthy that TMEM43-mediated current was inhibited dose-dependently by gradually lowering the extracellular pH from 8 to 5.5 with half-maximal inhibition at pH 6.495 (**Fig. 2i,j**), elucidating TMEM43 as a pH-sensitive channel. However, the change in the intracellular pH did not affect the channel current (**Extended Data Fig. 1g-i**). Altogether, these results define TMEM43 as an external pH-sensitive and nonselective cation channel.

**Fig. 1:**
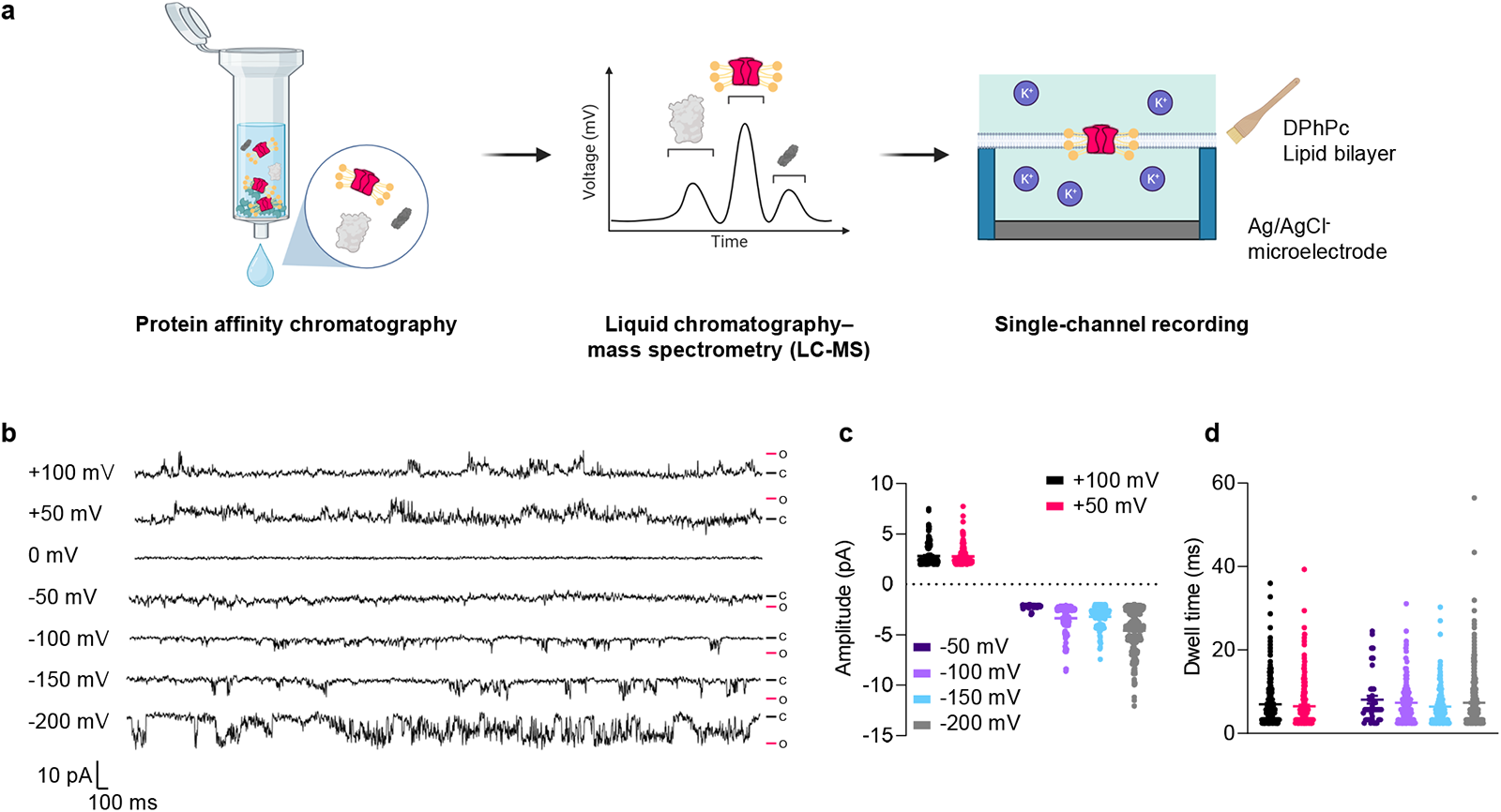
Single-channel activities of TMEM43. **a**, Illustration of experimental scheme. TMEM43 protein was purified in affinity column with Strep-tactin resin and isolated via LC-MS. The purified protein was used for parallel artificial lipid bilayer recordings. **b**, Stochastic channel gating of purified TMEM43 protein in different holding potentials. Closed states are indicated in the black bar with the letter C, and maximum open states are indicated in the pink bar with the letter O. (**c**, **d**) Distribution of amplitude (**c**) and dwell time (**d**) of open states.

**Fig. 2:**
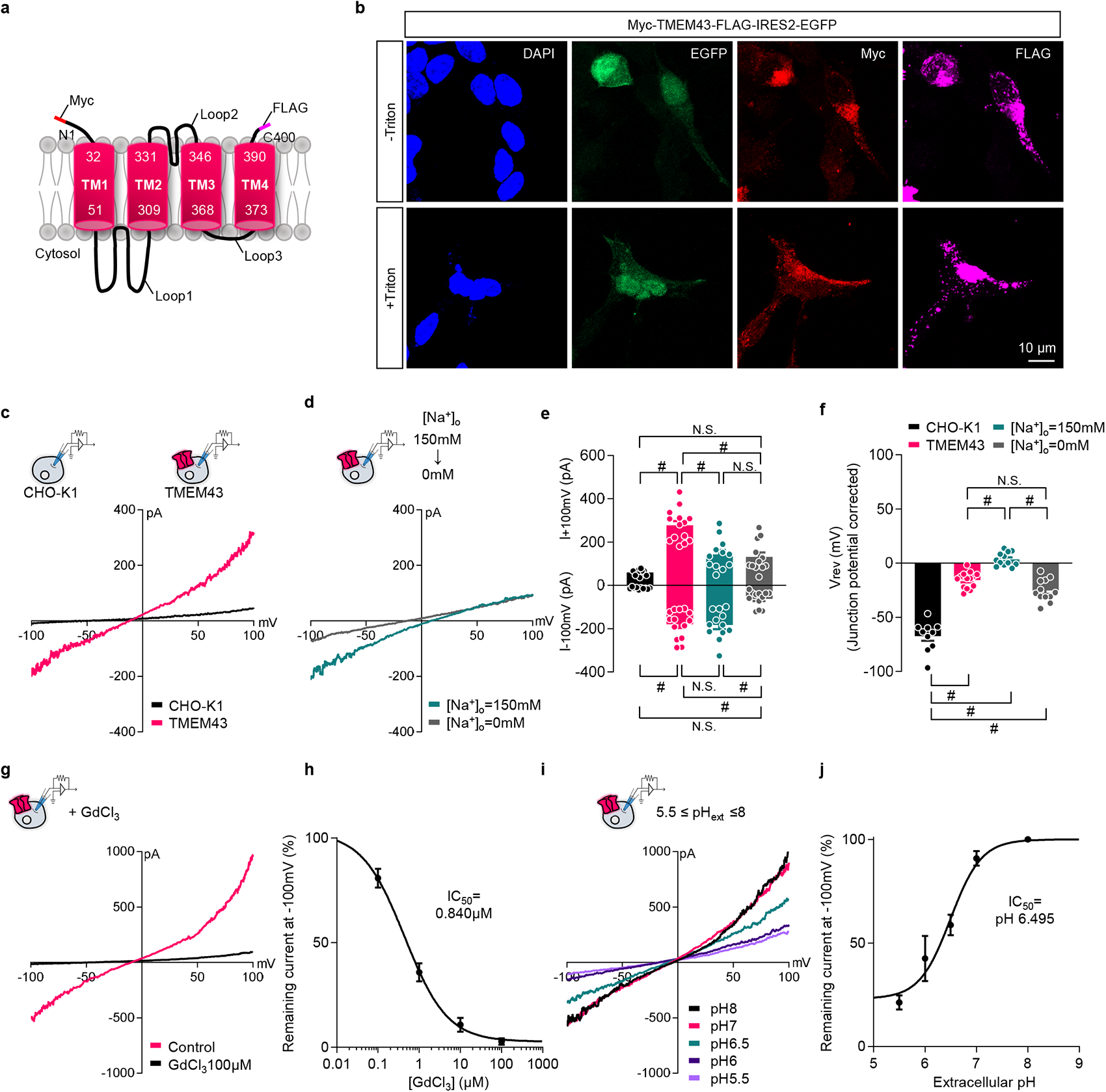
TMEM43 traffics to cell membrane and elicits pH-dependent cationic currents. **a**, Illustration of TMEM43 topology based on hydrophobicity analysis. Myc is tagged at the N-terminus, and FLAG is tagged at the C-terminus of TMEM43. **b,** Immunostaining result of TMEM43-expressing HEK293T cells with and without cell permeabilization. Cells were double stained with Myc and FLAG antibodies. **c**, Representative current traces from naïve (black) and *TMEM43* (Pink) transfected CHO-K1 cells. Current traces were elicited by 1-second ramps descending from +100 mV to -100 mV. Inset: a schematic diagram of patch-clamped cells. **d**, Representative current traces from *TMEM43* transfected cells in the presence (green) and absence (grey) of external Na^+^. The patch pipette is filled with N-methyl-D-glucamine (NMDG) gluconate, free of Na^+^ and K^+^. **e**, Summary bar graph of currents plotted at -100 mV (I_-100mV_) and +100 mV (I_+100mV_) for each condition in (**c**, **d**). **f**, Reversal potential (V_rev_) of respective conditions in (**c**, **d**) after junction potential correction. **g**, Representative whole-cell I-V curves from cells expressing *TMEM43* before (black) and after (pink) GdCl_3_ treatment (100 μM shown), measured in the same cell. **h**, Dose-response curve for GdCl_3_ blocking TMEM43 current. **i**, Representative current-voltage relationship in *TMEM43*-expressing cells with gradually decreasing external solution pH in the same cell. **j**, Dose-dependent inhibition of external solution acidic pH on TMEM43-mediated current. Data is presented as averaged % remaining currents ± SEM plotted at – 100 mV.

The TMEM43-mediated current increased dramatically in the hyperpolarizing potentials, negative to -100 mV (**Fig. 1b, Extended Data Fig. 1j**). This current-voltage (I/V) relationship from TMEM43-expressing cells highly resembled the current measured from inwardly rectifying potassium channel (Kir)-expressing cells^25,26^. Therefore, we checked if TMEM43 shares the characteristics of Kir channels. We found that treating barium (Ba^2+^), a selective blocker of Kir channels^27^, did not interrupt the TMEM43-induced large inward current at hyperpolarizing potentials (**Extended Data Fig. 1j,k**). This result indicates that the large inward currents at the hyperpolarizing potentials can be identified as a TMEM43-induced current independent of Kir channels. To determine the putative pore region of TMEM43 based on the membrane topology of TMEM43 (**Fig. 2a**), we performed the substituted-cysteine accessibility method (SCAM)^28^ using MTSES (Sodium (2-Sulfonatoethyl) methanethiosulfonate) as a reducing agent. MTSES binds strongly with the sulfhydryl groups of cysteine to block channel current if cysteine lies near the channel pore region^28,29^. The TMEM43 wild type (WT) contains only one cysteine residue (Cys354) at the transmembrane domain (TM)3. Surprisingly, treating 100 μM MTSES to TMEM43-expressing cell abated channel current by about 40 % (**Extended Data Fig. 2a,c**). This MTSES blockage disappeared when the endogenous cysteine at Cys354 was substituted to alanine p.(Cys354Ala) (**Extended Data Fig. 2b,c**), indicating that Cys354 is near or at the pore region of TMEM43. Furthermore, the reversal potential was significantly shifted after MTSES treatment in TMEM43 WT but not in TMEM43-p.(Cys354Ala) (**Extended Data Fig. 2d**), indicating that the mutation caused a change in ion selectivity. To further narrow down the channel pore region, we thoroughly examined the Loop2, which lies between TM2 and the cysteine-containing TM3. The deletion mutation of Loop2 (ΔLoop2) resulted in a complete loss of TMEM43-mediated current (**Extended Data Fig. 2e-g**). To check if the loss of current was due to a reduction in surface trafficking of TMEM43, we performed a biotinylation assay. We found that TMEM43-ΔLoop2 showed intact surface trafficking (**Extended Data Fig. 2h**), eliminating the possibility that TMEM43-ΔLoop2 fail membrane surface trafficking. Next, we substituted each of the ten amino acids at the Loop2 domain to cysteine in the TMEM43-p.(Cys354Ala) background to perform the SCAM experiment (**Extended Data Fig. 2i-k**). Averaged MTSES block percentage plot showed an ‘M’ shaped distribution with two maximal block percentages near two negatively charged aspartic acid residues of p.(Asp335Cys) and p.(Asp342Cys) with a middle dip at the kink-forming proline residue of p.(Pro338Cys) (**Extended Data Fig. 2k**). The SCAM results propose a putative model of the TMEM43 pore structure with two negatively charged residues per subunit to form a pore (**Extended Data Fig. 2k, inset**). These results strengthen the idea that TMEM43 is a *bona fide* ion channel with channel pore residues near the Loop2 domain, lining between TM2 and TM3.

The genetic variants of the human *TMEM43* gene have been highlighted in clinical studies. The *TMEM43*-p.(Arg372Ter) variant is reported to causes ANSD^5^ and the *TMEM43*-p.(Ser385Leu) variant is known to lead to ARVC in humans^3,4^. Therefore, we investigated whether TMEM43-mediated current is disturbed in the two *TMEM43* variants in the heterologous expression system. Whole-cell voltage clamp experiments revealed that *TMEM43*-p.(Arg372Ter) variant resulted in an abolishment of both inward and outward TMEM43-mediated current (**Extended Data Fig. 3a,b,d**), clearly documenting the pathogenic effect of the variant on the channel function. Furthermore, co-expression of *TMEM43*-WT and p.(Arg372Ter) variant (1:1) showed a significantly lower current than that of WT but comparable to that of mutant, especially at -100 mV, indicating that the pathogenic effect of the variant in TMEM43 is dominant-negative (**Extended Data Fig. 3c,d**). This result is compatible with the *TMEM43*-p.(Arg372Ter) autosomal dominant inheritance in humans as previously shown by linkage analysis and whole exome sequencing^5^. In contrast, *TMEM43*-p.(Ser358Leu) did not cause impairment in the nonselective cation current (**Extended Data Fig. 3e-g**), raising a possibility that the p.(Ser358Leu) and the p.(Arg372Ter) variants manifest different pathogenic mechanisms.

Gap junctions are intercellular channels, electrically and metabolically coupling nearby cells. Having found that TMEM43 can mediate channel currents, in addition to its interaction with gap junctions, we hypothesized that TMEM43 could be involved in mediating the transjunctional potentials. To investigate the electrical coupling property of the TMEM43 channels, we performed a dual-patch clamp recording from Naïve, TMEM43-, CX43-, and both TMEM43/Cx43-expressing HeLa cells (**Fig. 3a-c**). Although HeLa cells are coupling-deficient cells^30^, we observed bidirectional transjunctional potentials in receiver cells with current injections in sender cells after TMEM43 or Cx43 expression (**Fig. 3c and Extended Data Fig. 4a-c**). Remarkably, transjunctional potentials from TMEM43-expressing receiver cells were comparable to those of Cx43-expressing receiver cells (**Fig. 3c, d**). Moreover, we found that TMEM43 further enhances electrical couplings of Cx43 (**Fig. 3c, d**). These results demonstrate that TMEM43 mediates transjunctional potentials between two adjacent cells and facilitates gap junction couplings of connexins.

**Fig. 3:**
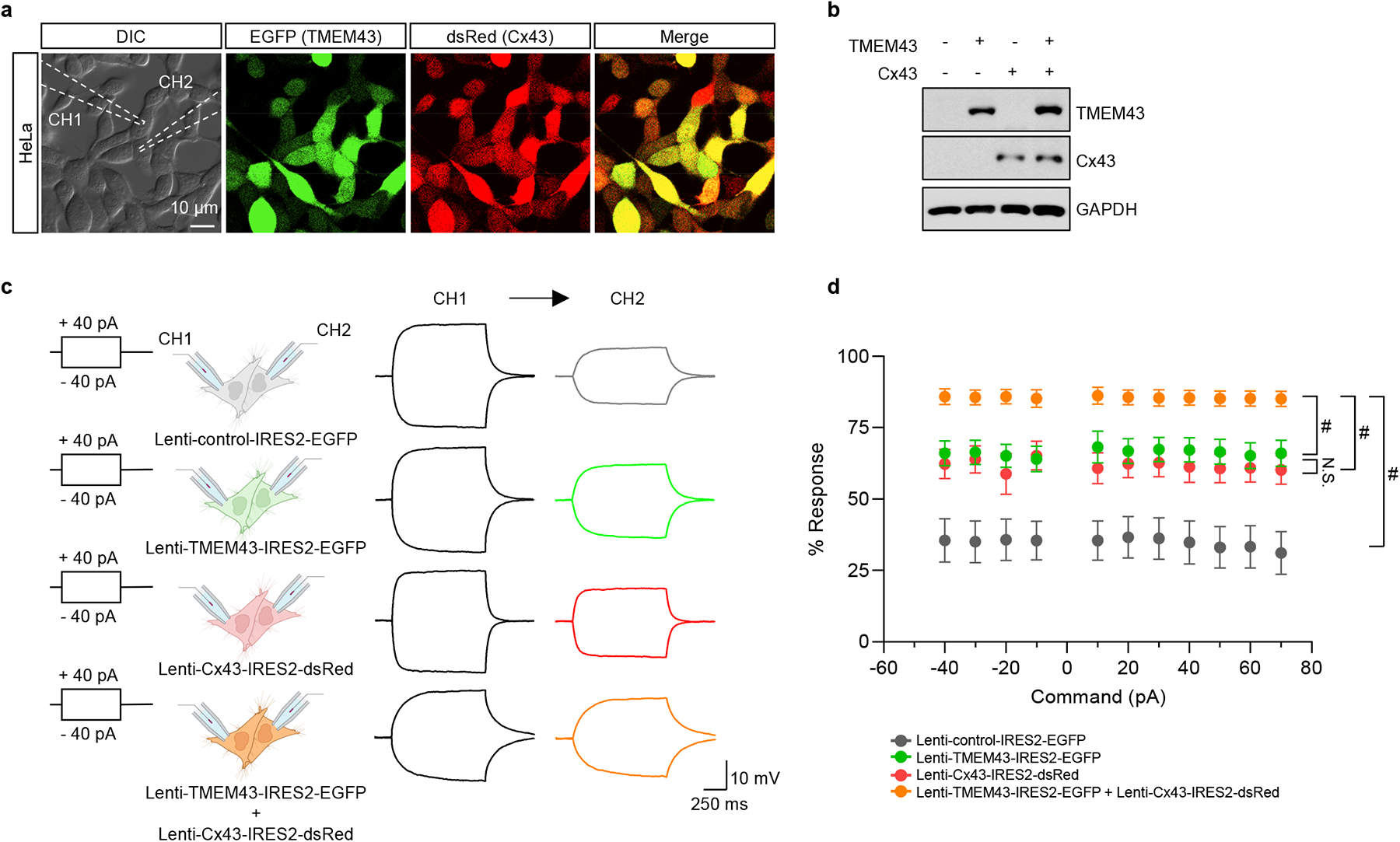
TMEM43 can mediate electrical coupling between cells. **a,** Images of HeLa cell culture infected with mixture of T*MEM43*-and *Cx43*-carrying lentivirus. Dotted white lines in DIC image outline two patch pipettes that have been used for dual-cell recording. CH1: channel 1, CH2: channel 2. **b**, Western blot result showing TMEM43 and Cx43 expression in the lentivirus infected HeLa cells. **c**, Experimental scheme of dual whole-cell recordings to measure electrical coupling between two contacting cells with lentivirus infections (left) and representative traces with +40 pA and -40 pA current injections (right). Current steps were given to sender cell via CH1 and voltage changes in receiver cell were measured via CH2. **d**, Averaged response percentage of transjunctional potentials in receiver cells measured form CH2 in control-(grey), TMEM43-(green), Cx43-(red), and TMEM43/Cx43 (orange)-expressing HeLa cells.

We moved on to examining the physiological role of TMEM43 in the brain. We dissected mouse brain in nine different brain regions including olfactory bulb, cerebellum, pons and medulla, midbrain, prefrontal cortex, striatum, hippocampus, diencephalon, and cerebral cortex. TMEM43 was detected in all examined brain areas (**Extended Data Fig. 5a**), indicative of its ubiquitous expression throughout the brain. To examine the function of TMEM43 in the brain, we took advantage of utilizing TMEM43 knockout (KO) mouse line (**Extended Data Fig. 5b-d**). The KO mice we used in this study has been designed to delete from Exon 2 to Exon 12 of *Tmem43* by CRISPR/Cas-mediated genome engineering, which covers 99% of the coding region including all four TM domains (**Extended Data Fig. 5b**, c). The efficiency of *Tmem43* gene deletion in the KO mice was confirmed by western blot assay (**Extended Data Fig. 5e**) and immunostaining (**Fig. 4a**). Considering high expression of connexin channels and their significant roles in hippocampus^11^, we examined TMEM43 expression in hippocampus. A bush-like TMEM43 expression was detected in hippocampal astrocytes, which was not detected in the TMEM43 KO (**Fig. 4a, b**). Of interest, astrocytes in TMEM43 KO were shown to possess thinned and elongated processes compared to WT astrocytes (**Fig. 4a**). Sholl analysis delineated that astrocyte morphology in TMEM43 KO is altered, with elongated filament length (**Fig. 4c-e**). We further examined TMEM43 expression in hippocampal astrocytes with the predominant connexin in the brain, Cx43^31,32^. The super resolution structured illumination microscopy (SIM) revealed TMEM43 and Cx43 expression in close proximity at hippocampal astrocytes, TMEM43 being closer to cytoskeleton (**Fig. 4f-h**). The co-IP assays demonstrated TMEM43 interaction with Cx43 in both in vitro heterologous expression system (**Extended data Fig. 6a**) and mouse brain lysates (**Extended data Fig. 6b**). Positive proximity ligation assays (PLA) signal was detected in hippocampal tissue with TMEM43 and Cx43 antibodies (**Extended data Fig. 6c**), strengthening the findings that TMEM43 is closely related to Cx43 in hippocampal astrocytes.

**Fig. 4:**
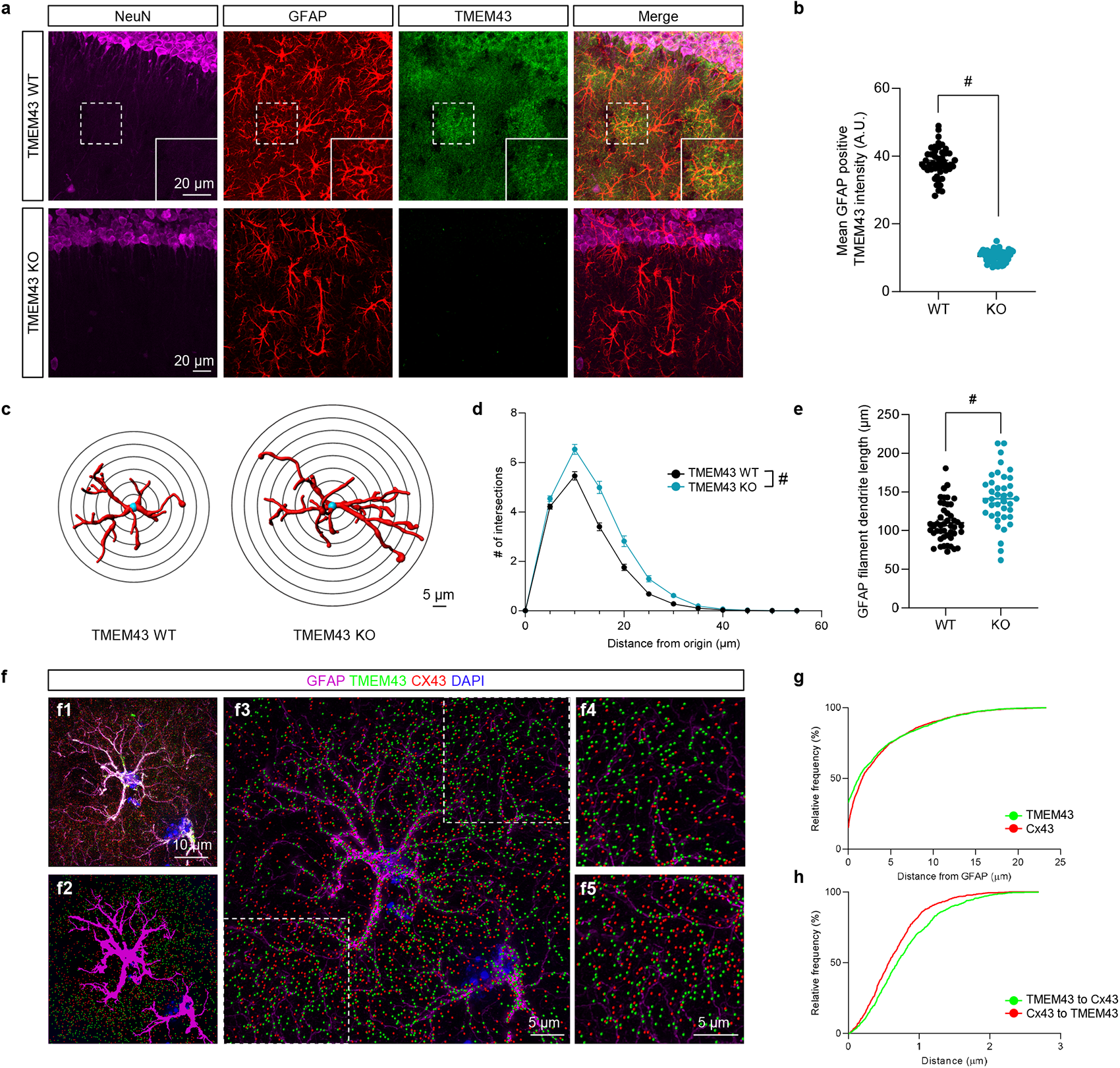
TMEM43 is expressed in astrocytes with Cx43 and lack of TMEM43 induce astrocyte hypertrophy. **a**, Spatiotemporal expression of TMEM43 (green) in hippocampal astrocytes (red). Immunoreactivity of TMEM43 was detected in the bush-like structure of astrocytes. Boxed area indicates magnified view of TMEM43 localization at GFAP-positive astrocyte. TMEM43 immunoreactivity was not found in KO tissue. **b**, Quantification of TMEM43 expression in GFAP-positive signals. **c**, Representative images of sholl analysis in astrocytes of TMEM43 WT (left) and KO (right). **d**, Number of intersections according to the distance from the astrocyte nucleus analyzed by sholl analysis. **e**, Averaged length sum of astrocyte branches of TMEM43 WT (black) and KO (blue). **f**, Representative images of SIM- and IMARIS-processed images. **f1**, SIM-processed image. **f2**, GFAP signal (magenta) reconstructed by ‘Surfaces’ function of IMARIS. **f3**, TMEM43 (green) and CX43 (red) reconstructed by ‘Spots’ function of IMARIS. **f4**, Magnified view of boxed area in the bottom left of (**f3**). **f5**, Magnified view of boxed area in the upper right of (**f3**). **g**, Cumulative frequency for distance of TMEM43 (green) and Cx43 (red) to reconstructed GFAP signal (**f2**). **h**, Cumulative frequency for distance of TMEM43 to Cx43 (green) and Cx43 to TMEM43 (red).

To study the functional role of TMEM43 in astrocytic gap junction coupling, we measured the number of coupled cells after patch clamping astrocytes with Alexa 488 (100 μM) dye-filled pipette and letting dye to diffuse for 20 min (**Fig. 5a**). The number of coupled cells were significantly reduced in TMEM43 KO compared to WT, which was further reduced with gap junction blocker carbenoxolone (CBX, 100 μM) treatment, indicating that the remaining portion of diffused cells in the KO comes from the presence of endogenous connexin channels (**Fig. 5b, c**). The vertical depth of coupled cells was also significantly decreased in TMEM43 KO compared to WT (**Fig. 5b, d**). However, we did not observe further reduction in vertical depth of coupled cells in the KO after CBX treatment, indicating that TMEM43 may be involved more in dye diffusion in vertical direction than the connexins (**Fig. 5b, d**). On the other hand, no significant differences in astrocytic passive conductance current or rectification index were found between WT and KO mice (**Extended data Fig. 7a-f**). Altogether, these results demonstrate that TMEM43 regulates gap junction function of cell-cell couplings in hippocampal astrocytes.

**Fig. 5:**
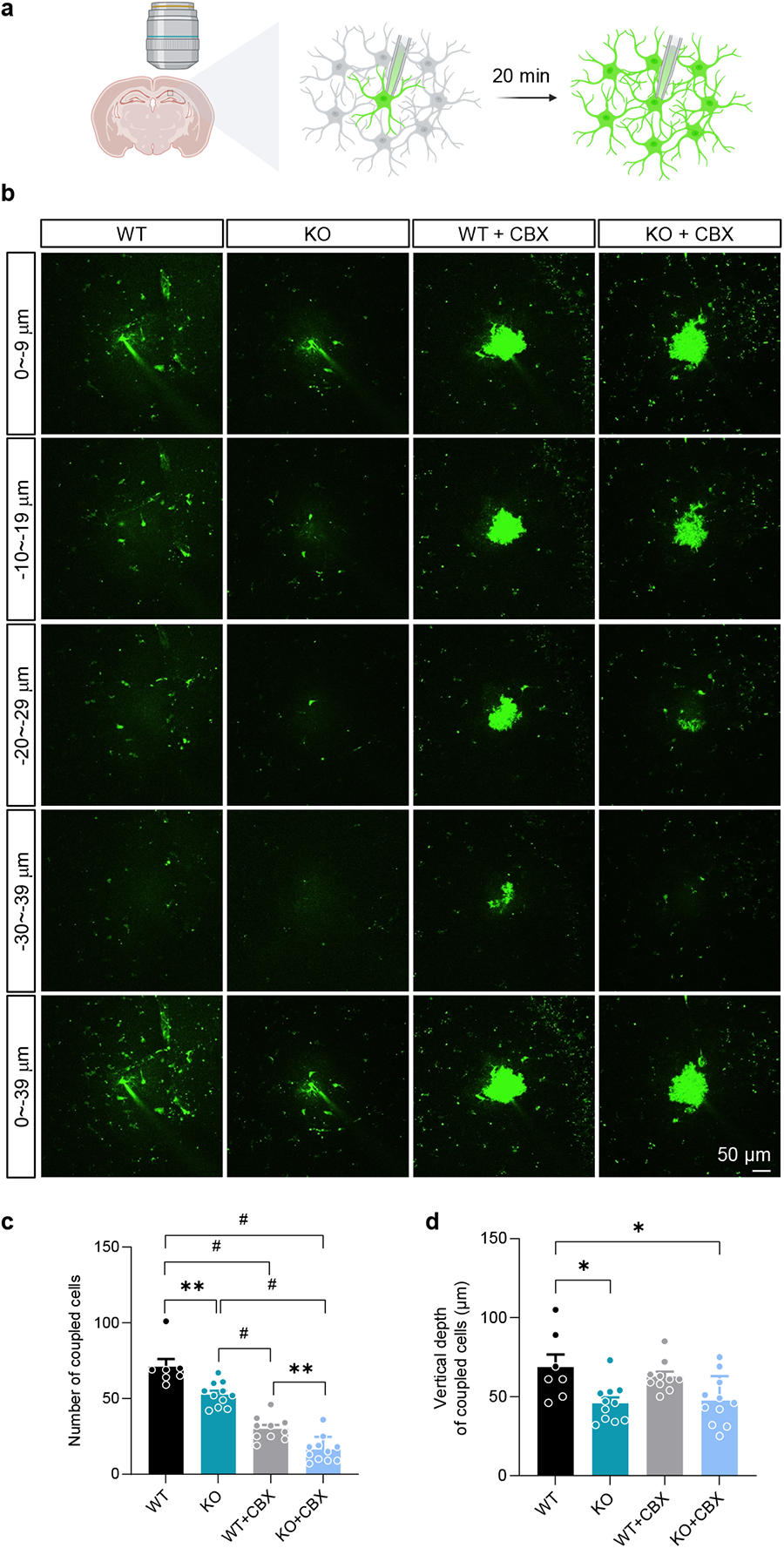
Gap junction coupling is disturbed in TMEM43 KO. **a,** Illustration of experimental scheme. Alexa 488 (100 μM) dye-loaded astrocytes were left to diffuse for 20 min. **b**, Representative images from two-photon microscope after astrocytic dye diffusion. **c**,**d**, Averaged number (**c**) and vertical depth of dye coupled astrocytes (**d**) in WT and KO mice, in presence and absence of CBX (100 μM) incubation.

We next investigated the impact of TMEM43 deletion on electrical properties of CA1 pyramidal neurons. Whole-cell patch clamp recordings in acute brain slices revealed that the rheobase current was significantly reduced in TMEM43 KO without changing input resistance (**Fig. 6a-d**). Significantly higher number of action potential (AP) firings with greater AP amplitude was observed in the KO (**Fig. 6e-h**). Although the WT and KO showed no significantly different pyramidal neuron resting membrane potentials (**Fig. 6i**), more current was generated in the KO at same given potential (**Fig. 6j**). These results indicate that intrinsic neuronal properties have been altered in the TMEM43 KO mice with enhanced neuronal excitability. We examined if this change in neuronal excitability disturbs AMPA/NMDA ratio. We found that Schaffer collateral stimulation-induced AMPA EPSC was significantly increased in the KO without changing NMDA EPSC, leading to disturbing the AMPA/NMDA ratio (**Fig. 6k-o**). Because NMDA decay tau was comparable between the two groups (**Fig. 6p**), we measured AMPA decay tau in presence of cyclothiazide (CTZ), a fast inhibitor of AMPA receptor desensitization. We found that AMPA decay tau was significantly increased in the KO (**Fig. 6q-s**), possibly explaining the increased cell excitability in the KO. Having found that the TMEM43 KO mouse elicit significantly reduced gap junction couplings, we hypothesized that the increase in AMPA decay tau in the KO could be attributable to the high K^+^ concentration in the extracellular space, caused by reduction in astrocyte K^+^ buffering. To test this idea, we infected astrocytes in stratum radiatum with a K^+^ sensor Green Indicator of K+ for Optical Imaging (GINKO1)^33^ (**Fig. 7a, b**). Schaffer collateral stimulation-induced K^+^ imaging disclosed that the K^+^ peak amplitude is decreased in TMEM43 KO with increase in both rise and decay tau (**Fig. 7c-f**), supporting the idea that K^+^ buffering is disturbed in the TMEM43 KO.

**Fig. 6:**
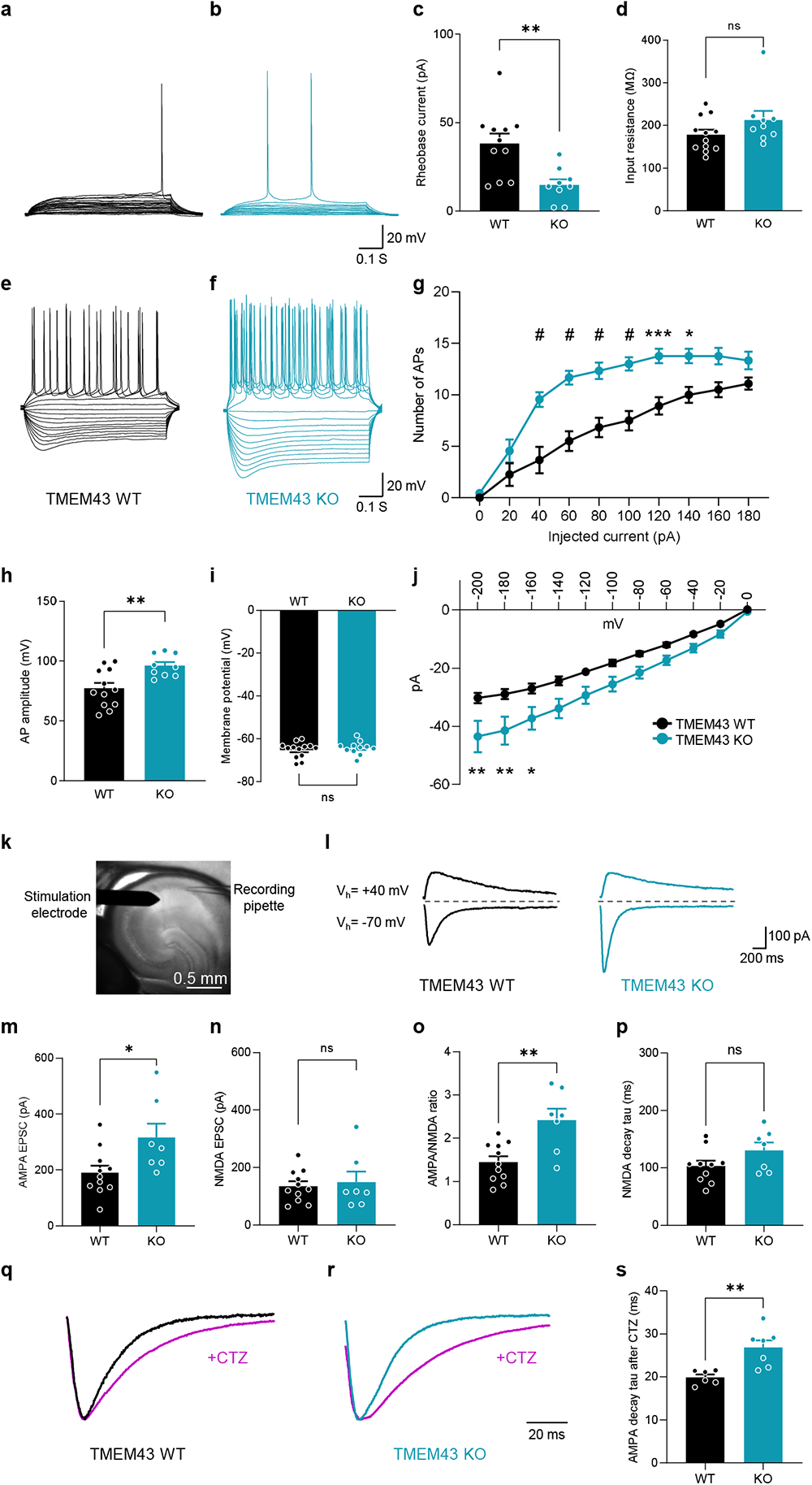
Hippocampal excitatory synaptic transmission is enhanced in TMEM43 KO. **a**,**b**, Representative traces of rheobase current from WT (**a**) and KO (**b**). **c**, Averaged rheobase current from WT and KO. **d**, Averaged input resistance from WT and KO. **e**,**f**, Representative traces of AP firing from WT (**e**) and KO (**f**). **g**, Number of APs in response to increasing current injection from WT and KO mice. **h**, Averaged AP amplitude from WT and KO. **i**, Averaged membrane potential from WT and KO. **j**, I-V relationship measured from WT and KO mice. **k**, Representative image showing hippocampal brain slice with stimulation electrode (left) and recording pipette (right). **l**, Representative AMPAR-and NMDAR-EPSC traces from WT and KO mice. AMPAR-mediated current was measured with holding potential of -70 mV and NMDAR-mediated current was measured with holding potential of +40 mV with CNQX. QX-314 was added in pipette internal solution and bicuculline was added in recording solution. **m**-**p**, Summary bar graphs of AMPA EPSC (**m**), NMDA EPSC (**n**), AMPA/NMDA ratio (**o**), and NMDA decay tau (**p**). **q**, Representative AMPAR-EPSC traces before (black) and after (purple) CTZ (100 μM) application in WT mice. **r**, Representative AMPAR-EPSC traces before (blue) and after (purple) CTZ (100 μM) application in KO mice. Traces were normalized by amplitude. **s**, Summary bar graph of AMPA decay tau after CTZ application.

**Fig. 7:**
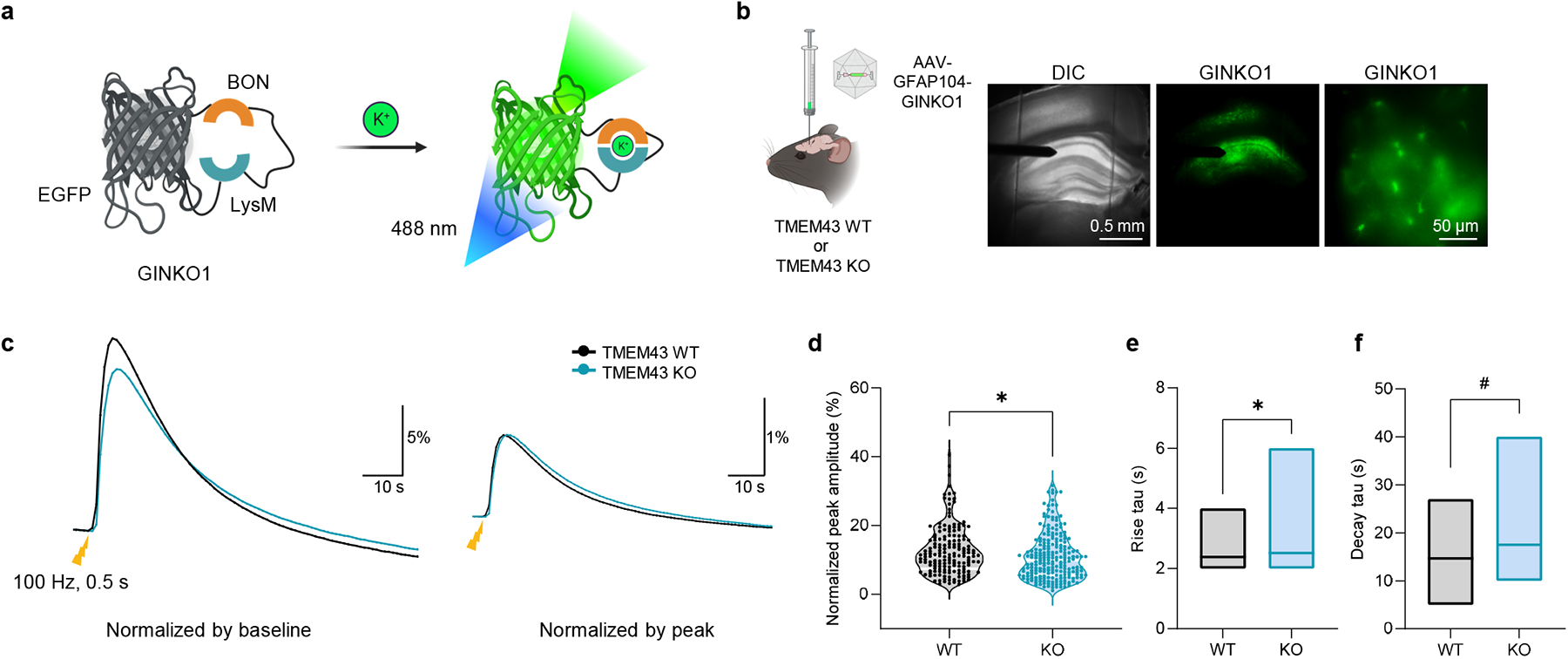
Potassium buffering is delayed in TMEM43 KO. **a**, Working mechanism of genetically encoded K^+^ indicator GINKO1. **b**, Experimental scheme for K^+^ imaging. AAV-GFAP104-GINKO1 was injected bilaterally in hippocampus stratum radiatum. Astrocytic GINKO1 expression was confirmed under 488 nm filter and electrode was placed in the Schaffer collateral pathway. **c**, Averaged GINKO1 response after stimulation in WT (black) and KO (blue). Left: traces normalized by baseline, right: traces normalized by peak. **d**, Normalized peak amplitude from (**c**). **e**,**f**, Floating bars of rise tau (**e**) and decay tau (**f**) from each group. Line in the box indicates mean.

Following the observations of astrocytic gap junction coupling and neuronal excitability alterations in hippocampus of TMEM43 KO, we tested weather hippocampus-dependent memory recall is disrupted in the KO. We performed a contexture fear conditioning test, measuring freezing time of a mouse, 24 hours after applying 6-foot shocks in random intervals (**Fig. 8a**). Acquisition memory was measured in the first day during habituation and shock applications. Time frozen increased gradually as more shocks were given, but we observed no noticeable differences between the WT and TMEM43 KO group (**Fig. 8b**). This result indicates that TMEM43 KO has intact learning acquisition memory. However, when we measured retention memory the following day, the freezing time of KO has significantly decreased compared to that of the WT (**Fig. 8c**). This result demonstrates that the retrieval memory had been impaired in the TMEM43 KO. To identify if long-term potentiation (LTP) underlies this impairment, we measured LTP in acute hippocampal brain slices from both groups with theta-burst stimulation (TBS) at the Schaffer collateral pathway (**Fig. 8d**). We found that the level of LTP is significantly decreased in the KO, explaining the underlying mechanism of the memory impairment (**Fig. 8e, f**). Next, we attempted to rescue the retention memory loss in the TMEM43 KO with TMEM43 overexpression. Mouse *Tmem43*-carrying AAV virus with GFAP104 promoter was delivered bilaterally to the hippocampus of KO (**Fig. 8g**). Astrocytic TMEM43 expression after viral infection was confirmed with immunostaining (**Fig. 8h**). The acquisition memory during learning was comparable in the KO and TMEM43-expressing KO (**Fig. 8i**). On the other hand, the retention memory was elevated in the TMEM43-expressing KO compared to vehicle-expressing KO (**Fig. 8j**), showing the sufficient evidence of TMEM43 contributing to hippocampal memory retrieval. Basal locomotion activity (**Extended data Fig. 8a-c**) or short-term memory (**Extended data Fig. 8e-j**) was not altered in the KO. Altogether, we demonstrate the critical role of TMEM43 in gap junction networks to regulate long term memory retrieval.

**Fig. 8:**
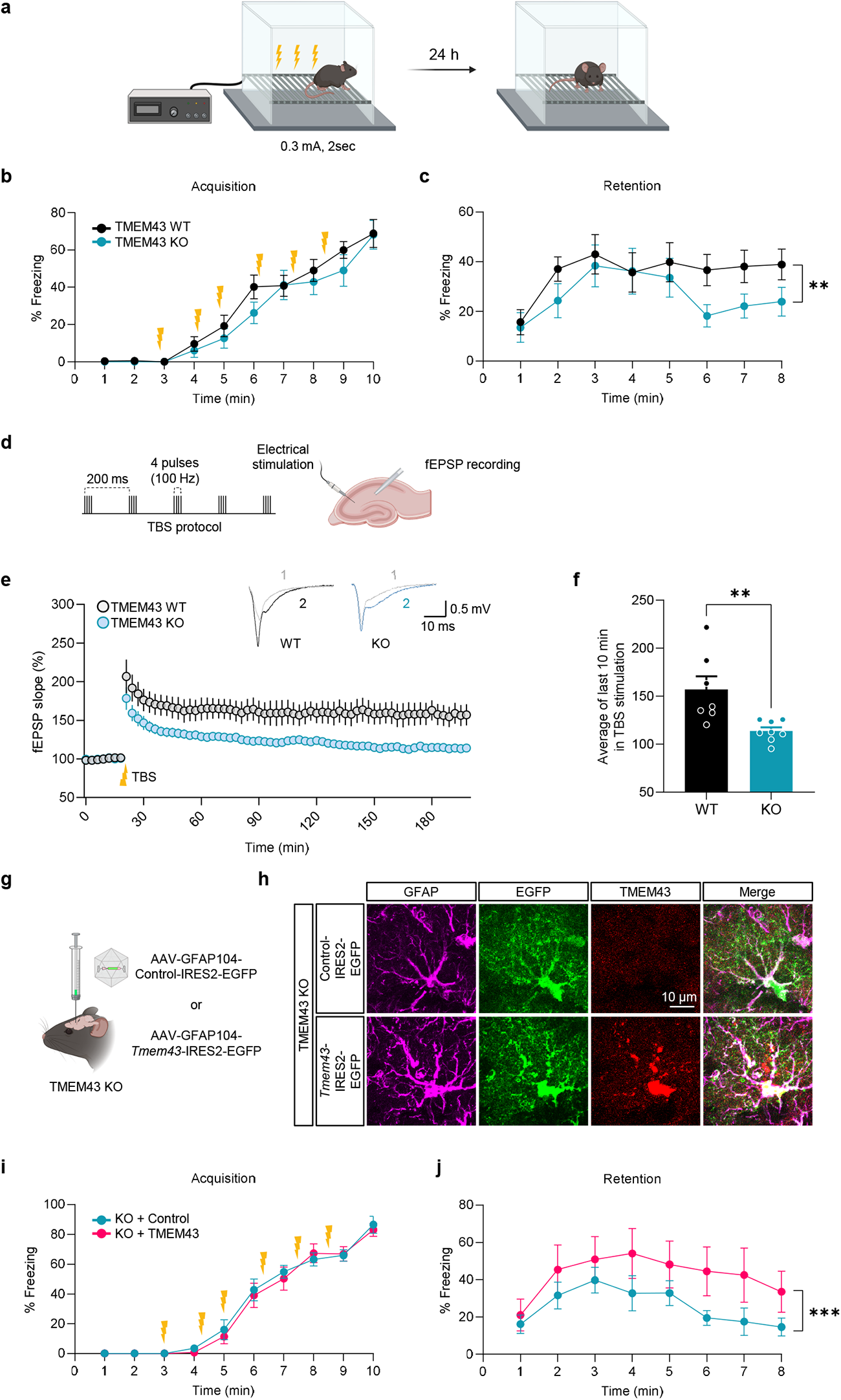
TMEM43 KO shows reduced memory retention which is rescued by TMEM43 overexpression. **a,** Illustration of experimental scheme for contexture fear conditioning test. 24-hour memory was tested. **b**,**c**, Averaged percentage of freezing time during acquisition session (**b**) and retention session (**c**) in WT (black) and KO (blue). Yellow symbols represent delivery of electric shock (0.3 mA, 2 sec). **d**, Schematic diagram of fEPSP recording. TBS protocol consists of short train stimulation that contains 4-pulses burst (100 Hz) repeated 5 times in 200 ms intervals. **e**, TBS-induced LTP in WT (black) and KO (blue) under fEPSP recording. Inset 1 trace; the representative trace before electrical stimulation. Inset 2 trace; the representative trace after electrical stimulation. **f**, Summary graph of TBS-induced LTP in WT (black) and KO (blue). **g**, Illustration of experimental scheme for animal model. Either control virus or *Tmem43*-carrying virus was injected to the hippocampus of TMEM43 KO mice. **h**, Immunostaining results validating viral expression and TMEM43 overexpression. **i**,**j**, Averaged percentage of freezing time during acquisition session (**i**) and retention session (**j**) in KO with control virus (blue) and KO with TMEM43 overexpression (pink). Yellow symbols represent delivery of electric shock (0.3 mA, 2 sec).

## Discussion

TMEM43 was initially identified in the inner nuclear membrane ^8^ and *in silico* as one of the 368 transmembrane proteins with unknown function (TMEM) ^34^. A few unknown TMEMs have been identified as ion channels, including ANO1 (TMEM16A) as a Ca^2+^-activated anion channel ^35^ and TENTONIN3 (TMEM150C) as a mechanosensitive cation channel ^36^. Although pathological functions of TMEM43 have been implicated in many human diseases, the characterization of the TMEM43 protein itself was missing in the field. It has been demonstrated that disruption of TMEM43 protein in the cochlea by gene-silencing, introducing the genetic variant *TMEM43*-p.(Arg372Ter), and interrupting TMEM43-interacting proteins such as Cx26, Cx30, and TASK-1, all resulted in the loss of K^+^ conductance current in the cochlear glia-like supporting cells ^5,9^. We got insight from the previous studies and hypothesized that TMEM43 may encode a new class of ion channels. In this study, we utilized purified TMEM43 protein and heterologous gene expression system combined with intensive electrophysiology and identified TMEM43 as an acid-sensing cation channel. Based on the predicted TMEM43 topology, we characterized the putative channel pore domain to be including the Loop2 domain. It is notable that TMEM43-expressing cells conduct transjuctional potentials between two adjacent cells, highly resembling the connexin channels, and further enhancing electrical couplings of the connexins. This result suggests TMEM43 as a secondary protein to efficiently modulate gap junction couplings between cells. We were aware of different ion conductance in *TMEM43*-p.(Arg372Ter) and *TMEM43*-p.(Arg358Leu)-expressing cells. This difference likely comes from the absence or presence of the TM4 domain, as the nonsense p.(Arg372Ter) variant results in truncation of the TM4, while the missense p.(Arg358Leu) variant still retains TM4. Lack of the TM4 may decompose TMEM43 oligomerization, disrupt secondary protein binding or missing in ion-conducting passage, which can all lead to loss of channel currents. Future research to visualize TMEM43 protein structure, such as by the cryo-EM-based protein structure reconstruction, would provide answers to the unresolved questions including identification of channel pore domain, pH-sensing domain and protein-protein interaction domain.

We took advantage of TMEM43 null KO mice in understanding the physiological role of TMEM43 in the brain. TMEM43 expression was detected in bush-like structures of hippocampal astrocytes along with Cx43. We demonstrated that TMEM43 is actively involved in mediating astrocytic gap junction couplings by measuring astrocyte dye-couplings. Intriguingly, TMEM43 was shown to be involved more in regulating gap junction couplings in ventral directions than the connexin channels do. The action of modulatory or secondary proteins in shaping the directions of gap junction couplings is a compelling subject to be studied in time to come. A unique astrocytic morphology was observed in the TMEM43 KO, with longer and thinner processes. We speculate that astrocytes with decreased electrical couplings may attempt to compensate loss of gap junction couplings by elongating their processes to increase physical contacts. The astrocytic morphology in TMEM43 KO clearly differs from astrocyte-specific Cx30 and Cx43 deficient mice, where the astrocytes display morphology of reactiveness, entailing inflammations^14^. The detailed mechanism should be investigated in future. Extensive whole-cell patch clamp recordings revealed that CA1 pyramidal neurons in TMEM43 KO share the hyperexcited neuronal properties observed in the Cx30 and Cx43 double-KO mice^13^. Neurons in both TMEM43 and connexin-lacking mouse lines elicit enhanced AP firings, reduced rheobase current, and increased AMPA EPSC. Astrocytes adopt more negative membrane potential than neurons by expressing potassium channels for mediating leak passive conductance current and facilitating potassium buffering^37,38^. We noticed substantial reduction in astrocytic buffering system and gap junction-mediated diffusions in the TMEM43 KO, permitting neurons to depolarize^39^. The hyperexcited neurons in TMEM43 KO eventually led to disruption in AMPA/NMDA ratio and LTP induction, explaining the reduced memory retention in the TMEM43 KO. On the other hand, the short-term memory and memory acquisition of TMEM43 KO were comparable to the WT, highlighting the role of TMEM43 in enhancing and fine-tuning gap junction networks in the brain. This study will expedite further studies in exploring gap junction coupling regulations and finding therapeutic targets for TMEM43 and connexin-related pathology.

## Methods

### Animals

Both male and female C57BL/6J-*Tmem43*^em1C^/Cya knockout (Cyagen, #S-KO-14294) mice were used at 6-16 weeks of age. All animals were housed in a 12-h/12-h light-dark cycle in a specific-pathogen-free facility with controlled temperature and humidity and had free access to food and water. Mice were group housed, but no more than five per cage. All experimental procedures were conducted according to protocols approved by the directives of the Institutional Animal Care and Use Committee of IBS (Daejeon, Republic of Korea).

### Plasmids

*TMEM43* Human Tagged ORF Clone (NM_024334.2) was purchased from OriGene (RC200998) and cloned into CMV-MCS-IRES2-EGFP vector using BglII/XmaI sites. *TMEM43* variants, including *TMEM43*-p.(Arg372Ter), *TMEM43*-(p.Ser358Leu), and site-directed mutations were obtained by performing oligonucleotide-directed mutagenesis using the EZchange site-directed mutagenesis kit (EZ004S, Enzynomics) and cloned into CMV-MCS-IRES2-EGFP or CMV-MCS-IRES2-tdTomato vector using BglII/XmaI sites. KCNK3 (Myc-DDK-tagged) (TASK-1) (NM_002246.3) was purchased from OriGene (RC215155) and cloned into IRES2 vector using BglII/XhoI sites, and K_ir_4.1 (NM_031602) was purchased from addgene (52874) and cloned into IRES2 vector using XhoI/SalI sites.

### Heterologous expression of TMEM43 in mammalian cell lines

Human embryonic kidney (HEK) 293T cells and Chinese hamster ovary (CHO)-K1 cells were each purchased from ATCC (CRL-3216) and the Korean Cell Line Bank (10061, Seoul National University), respectively. All cell lines have been tested for mycoplasma contamination. Cell lines were cultured in DMEM (10-013, Corning) for HEK293T cells and Ham’s F12 medium (21127-022, Gibco) for CHO-K1 cells, which were supplemented with 10 % heat-inactivated fetal bovine serum (10082-147, Gibco) and 10,000 units/ml penicillin-streptomycin (15140-122, Gibco) at 37 °C in a humidified atmosphere of 95 % air and 5 % CO_2_. Transfection of expression vectors was performed with Effectene Transfection Reagent (Effectene, 301425, Qiagen), according to the manufacturer’s protocol. One day before performing whole cell patch experiments, CHO-K1 cells were transfected with plasmid DNA 1 μg per 35 mm dish. The ratio of DNA to Effectene Reagent is 1:10.

### Immunocytofluorescence on heterologous system

For immunocytochemistry, pCMV-*TMEM43*-IRES2-EGFP were transfected into HEK293T cells one day before staining. Cells were fixed in 4 % paraformaldehyde for 10 minutes at room temperature and washed 3 times with PBS. The permeabilized group contained 0.3 % Triton X-100 in the blocking solution with 2 % goat serum and 2% donkey serum, but the impermeabilized group excluded Triton X-100 in the blocking solution. Cells were incubated with mouse anti-Myc (1:500, 2276, cell signaling) and rabbit anti-FLAG (1:500, 2368, cell signaling) at 4°C overnight. After washing, donkey anti-mouse Alexa 594 (1:500, Jackson, 715-585-150) and donkey anti-rabbit Alexa 647 (1:500, Jackson, 711-605-152) were added and incubated for 2 hours at room temperature. The cells were washed 3 times, mounted on slide glasses, and observed under a Zeiss confocal microscope (LSM900).

### Protein purification

TMEM43-EGFP-thrombin-twinstrep (1 µg) was expressed in Expi293 cell line (3X10^6^ cells/ml, 200 ml in 1 L flask) using ExpiFectamine™ 293 Transfection Kit (Gibco, A14524). After 48 hours, cells were lysed using a sonicator, and cell debris was removed by 4000 rpm, 30 min, 4°C centrifugation. Then, 150000 g, 1 hour, 4°C ultra-centrifugation was done to separate membrane proteins. The membrane pellet was mechanically homogenized and solubilized for 2 hours at 4°C in a buffer that contained 250 mM NaCl, 50 mM HEPES (pH 7.5) in 1% n-Dodecyl-β-D-Maltoside Lauryl Maltoside Dodecyl 4-O-α-D-Glucopyranosyl-β-D-Glucopyranoside (DDM) (Antrace, 69227-93-6) and 0.1% Cholesteryl Hemisuccinate Tris Salt (CHS) (Antrace, 102601-49-0). Insoluble material was removed by ultra-centrifugation (150000 g, 1 hour, 4°C). Finally, TMEM43 protein was filtered using Strep-tactin resin (IBA, 2-1201-025) and eluted with 0.02% DDM and 10 mM D-biotin (IBA, 2-1000-005).

### Single-channel recordings

The Orbit mini apparatus (Nanion, Germany, horizontal planar lipid bilayer system) was used to record single-channel activities. The Orbit mini is a miniaturized bilayer workstation that simultaneously enables recording from four artificial lipid bilayers. The recordings were done in a symmetrical 140 mM KCl (pH 7.2) solution at room temperature. DPhPC (1 mg/ml) dissolved in Octane was used to paint a bilayer over the four 100 µm diameter wells in a Meca4 chip (Nanion). Purified TMEM43 protein in DDM detergent (0.02 %) was loaded into the bath solution after forming the bilayers.

### Cell surface biotinylation, co-immunoprecipitation, and western blot

For biotinylation, plasmid vectors were transfected into HEK293T cells 1 day before the experiment day. Transfected cells were washed three times with PBS, and cell surface-expressed proteins were biotinylated in PBS containing Ez-link sulfo-NHS-LC-Biotin (21335, Thermo) for 30 minutes. After biotinylation, cells were washed with quenching buffer (100 mM glycine in PBS) to remove excess biotin and washed three times with PBS. The cells were then lysed and incubated with high-capacity NeutrAvidin-Agarose Resin (29204, Thermo). After three washes with lysis buffer, bound proteins are eluted by the SDS sample buffer and subjected to western blot analysis. Rabbit anti-TMEM43 polyclonal (1:250, NBP1-84132, Novus) and rabbit anti-β-Actin (1:1000, 4970L, Cell signaling) were used as primary antibodies, and donkey anti-rabbit HRP (NA9340, Amersham) was used as a secondary antibody. For co-immunoprecipitation, cell lysates were prepared in a buffer containing 50 mM Tris-HCl (pH 7.5), 150 mM NaCl, 1 % NP-40, 10 mM NaF, and a protease and phosphatase inhibitor cocktail. Equal amounts of precleared cell lysates were incubated with rabbit anti-TMEM43 polyclonal (1 μg, NBP1-84132, Novus) overnight at 4°C. Protein A/G-Agarose beads (Thermo Fisher Scientific) were added to the mixtures and further incubated for 2 hours, followed by a wash with lysis buffer. Bound proteins were eluted from the beads with SDS-PAGE sample buffer, and western blotting was performed with mouse anti-TASK-1(KCNK3) (1:250, NBP2-42202, Novus) or rabbit anti-K_ir_4.1 (1:250, APC-035, Alomone Labs).

### Structured illumination microscopy (SIM)

The super-resolution images were obtained using the Elyra 7 super-resolution microscope and processed using ZEISS Zen 3.0 software. The laser intensities are as follows: 405nm, 17%; 488 nm, 7%; 561 nm, 7%; 642 nm, 7%.

### Electrophysiological recording in CHO-K1 cells

Current-voltage (I–V) curves were recorded from CHO-K1 cells one day after plasmid DNA transfection. I–V curves were established by applying 1-second duration voltage ramps descending from +100 mV to -100 mV, with -60 mV holding potential. Transfected CHO-K1 cells were distinguished by fluorescence using an Olympus IX71 inverted microscope. Recording pipettes were fabricated from borosilicate glass capillaries (TW150F-4, World Precision Instruments) using a P-97 Flaming/Brown micropipette puller (Sutter Instruments). The standard bath solution contained (in mM): 150 NaCl, 3 KCl, 2 CaCl2, 2 MgCl2, 10 HEPES, and 5.5 D-glucose (pH 7.4 was adjusted with NaOH). For measuring potassium currents, recording electrodes (5–8 MΩ) were filled with (mM): 126 KOH, 126 Gluconate, 5 HEPES, 0.5 MgCl_2_, and 10 BAPTA (pH adjusted to 7.3 with KOH). To measure sodium currents, recording electrodes (5–8 MΩ) were filled with (mM): 126 NMDG, 126 Gluconate, 5 HEPES, 0.5 MgCl_2_, and 10 BAPTA. Then, the bath solution was shifted from standard buffer to NMDG buffer (in mM: 120 NMDG-Cl, 10 HEPES, 3 KCl, 2 CaCl_2_, 2 MgCl_2_, and 5.5 glucose, pH 7.3). For measuring cesium current and calcium-dependent current, recording electrodes (5–8 MΩ) were filled with (mM): 106 CsCl, 20

TEA-Cl, 5 HEPES, 0.5 MgCl_2_, 10 BAPTA, and 0 and 9.95 CaCl_2_ to make 0 μM and 10 μM Ca^2+^ (pH adjusted to 7.3 with CsOH). For measuring chloride currents, recording electrodes (5–8 MΩ) were filled with (mM): 20 TEA-Cl, 5 HEPES, 0.5 MgCl_2_, 10 BAPTA, and each 106 gluconate and 106 CsCl to make 1 mM Cl^-^ and 127 mM Cl^-^ (pH adjusted to 7.3 with CsOH). To measure the external pH-dependent current, HEPES buffer was made and adjusted initially to pH 5 with NaOH, and pH was increased serially by adding NaOH. To measure the internal pH-dependent current, the pH of the K-gluconate internal solution was serially adjusted with KOH from pH 5 to 8. Whole-cell membrane currents were amplified by the Axopatch 200A, and data acquisition was controlled by pCLAMP 10.2 software (Molecular Devices). The Digidata 1322A interface was used to convert digital-analog signals between the amplifier and the computer. Data were sampled at 10 kHz and filtered at 2 kHz. Cell membrane capacitance was measured using the ’membrane test’ protocol built into pCLAMP. All experiments were conducted at a room temperature of 20-22°C. All I-V curves were constructed without liquid junction potential correction. However, the reversal potentials were calculated by subtracting the measured liquid junction potential (-14 mV) from the observed values.

### Electrophysiological recording in HeLa cells

Electrical couplings were performed from HeLa cells two or 3 days after virus infection. Transfected CHO-K1 cells were distinguished by fluorescence using a Nikon A1R Confocal/inverted microscope. Electrical couplings were obtained from receiver cells in I=0 mode, by applying 1-second duration current steps (+70 pA to -40 pA) in sender cells at resting membrane potential with current clamp mode. Recording pipettes were fabricated from borosilicate glass capillaries (TW150F-4, World Precision Instruments) using a P-97 Flaming/Brown micropipette puller (Sutter Instruments). The standard bath solution contained (in mM): 150 NaCl, 3 KCl, 2 CaCl2, 2 MgCl2, 10 HEPES, and 5.5 D-glucose (pH 7.4 was adjusted with NaOH). For measuring potassium currents, recording electrodes (5–8 MΩ) were filled with (mM): 126 K-gluconate, 5 HEPES, 0.5 MgCl_2_, and 10 BAPTA (pH adjusted to 7.3 with KOH). Whole-cell membrane currents were amplified by the Multiclamp 700B, and data acquisition was controlled by pCLAMP 11.2 software (Molecular Devices). The Digidata 1550B interface was used to convert digital-analog signals between the amplifier and the computer. Data were sampled at 10 kHz and filtered at 2 kHz. All experiments were conducted at a room temperature of 20-22°C.

### Evoked AMAP receptors-mediated currents

AMPAR-EPSCs were isolated with 200 μM DL-AP5 and 20 μM bicuculline methiodide to block NMDA and GABAA receptors, and 5mM QX-314 was added to the internal solution to block voltage-gated Na+ channels. EPSCs were evoked using a monopolar platinum stimulator applying a 100 ms duration constant current stimulation to the Schaffer-collaterals in the hippocampus. Cyclothiazide (100 μM) was applied to the external solution for 15 minutes to block the AMPAR desensitization.

### Astrocytic dye-diffusion

Hippocampal horizontal acute slices (300 μM thickness) were prepared from TMEM 43 WT and KO mice aged around 10 to 14 weeks old and maintained at room temperature in a submerged chamber with extracellular artificial CSF (aCSF) solution (in mM): 126 NaCl, 24 NaHCO_3_, 1 NaH2PO4, 2.5 KCl, 2.5 CaCl_2_, 2 MgCl_2_, and 10 d-(+)-glucose (pH 7.4) All the solutions were bubbled with 95% O_2_ and 5% CO_2_, as previously described (brain paper). Slices were incubated at room temperature for at least 1 hour before recording with gas bubbling. Slices were transferred to a recording chamber that was continuously perfused with aCSF using a peristaltic pump (Miniplus 3, Gilson) at a rate of 2–3 ml/min. Whole-cell configurations were conducted from astrocytes in the Schaffer Collateral at holding potential of −80 mV. Pipette resistance was typically 8 to 10 MΩ. The pipette was filled with the internal solution (in mM): 80 K-gluconate, 60 KCl, 10 HEPES, 5 EGTA, 2 Mg-adenosine triphosphate, 0.5 Na_2_-guanosine triphosphate, 0.1 Alexa 488, pH adjusted to 7.3 to 7.4 with KOH (osmolarity, 275 to 285 mOsm)]. In order to detect dye diffusion among astrocytes, we used the Nikon A1R Multiphoton (MP) microscope, optically linked to Spectra-Physics Mai-Tai IR variable wavelength pulse laser (690-1040nM). Fluorescence images were obtained 60–120 optical sections in the Alexa emission channel (540LP/700SP filter; λ2px = 830 nm), collected in image frame mode (512 pixels × 512 pixels, eight-bit) at 1 μm step at least 20 min after a whole-cell configuration.

### Chemicals

Gadolinium chloride (GdCl_3_) was purchased from Sigma-Aldrich (G7532); Barium chloride (BaCl2) was purchased from Sigma-Aldrich (B0750); Sodium (2-Sulfonatoethyl) methanethiosulfonate (MTSES) was purchased from Toronto Research Chemicals (S672000).

### Behavior tests

#### Open field test

Mice were placed into an open field chamber (40 cm × 40 cm × 40 cm), and allowed to freely explore the area for 10 minutes. The center is defined as an area in the middle of the chamber (20 cm x 20 cm). An automated tracking system (EthoVision) was used to monitor and analyze the animal’s locomotion.

#### Novel place recognition test

On the next day of the open field test, mice were placed into the same open field chamber with two identical objects positioned in the first and the second quadrant of the arena and allowed to freely explore the objects for 10 min. Then, the mice were put back in the home cage. After 1 hour, mice were placed into the chamber again to explore the objects for 10 min, but one of the two objects had been relocated into the opposite empty quadrant of the arena. The exploration behavior of each mouse was video recorded and analyzed manually by an experimenter. The object exploration was defined as direct contact of the nose with the object. Climbing on top of the object was not considered as exploration.

#### Y Maze Spontaneous Alternation Test

A Y-shaped maze with three white, opaque plastic arms at a 120° angle from each other was used. A mouse was firstly placed in the maze, facing towards the center. The animal was then left to freely explore the three arms for 8 min. Arm entry was defined as mouse passing the first quarter of a arm.

#### Contextual fear conditioning

The chamber (20cm x 20cm x 30cm) with a stainless-steel floor and a camera mounted on its ceiling (Coulbourn Instruments, model: H10-11M-TC) was located in a sound-proof box. On the first day, mice were allowed to explore the chamber freely for 3 min, and received six foot-shocks in random time intervals (at 180 sec, 250 sec, 300 sec, 380 sec, 440 sec, and 510 sec) for 2 seconds in 0.3 mA. Twenty-four hours after the training, the mice were placed in the chamber for 10 min and their behavioral responses were videotaped. Freezing response, defined as an absence of any movement except breathing for >1 sec, was measured and analyzed using FreezeFrame 3.0 software (Coulbourn Instruments).

#### Data analysis and statistical analysis

Off-line analysis was carried out using Clampfit version 11.2 and GraphPad Prism version 9 software. When comparing two samples, the significance of data was assessed by Student’s two-tailed unpaired t-test when samples showed normal distribution and assessed by the Mann-Whitney test when samples did not pass the normality test. Samples that passed the normality test but not the equal variance test were assessed with Welch’s correction. Comparing more than 2 samples were analyzed using one-way ANOVA with Tukey’s post-hoc test when data passed the normality test and Kruskal-Wallis test with Dunn’s post-hoc test when data did not pass the normality test. Significance levels were given as: N.S. *P*>0.05, \**P*<0.05, \*\**P*<0.01, \*\*\**P*<0.001 and #*P*<0.0001 in all figures.

## Acknowledgments

This research was supported by Institute for Basic Science (Grant/Award Number: IBS-R001-D2) to C. Justin Lee.

---

**Extended Data Fig. 1:**
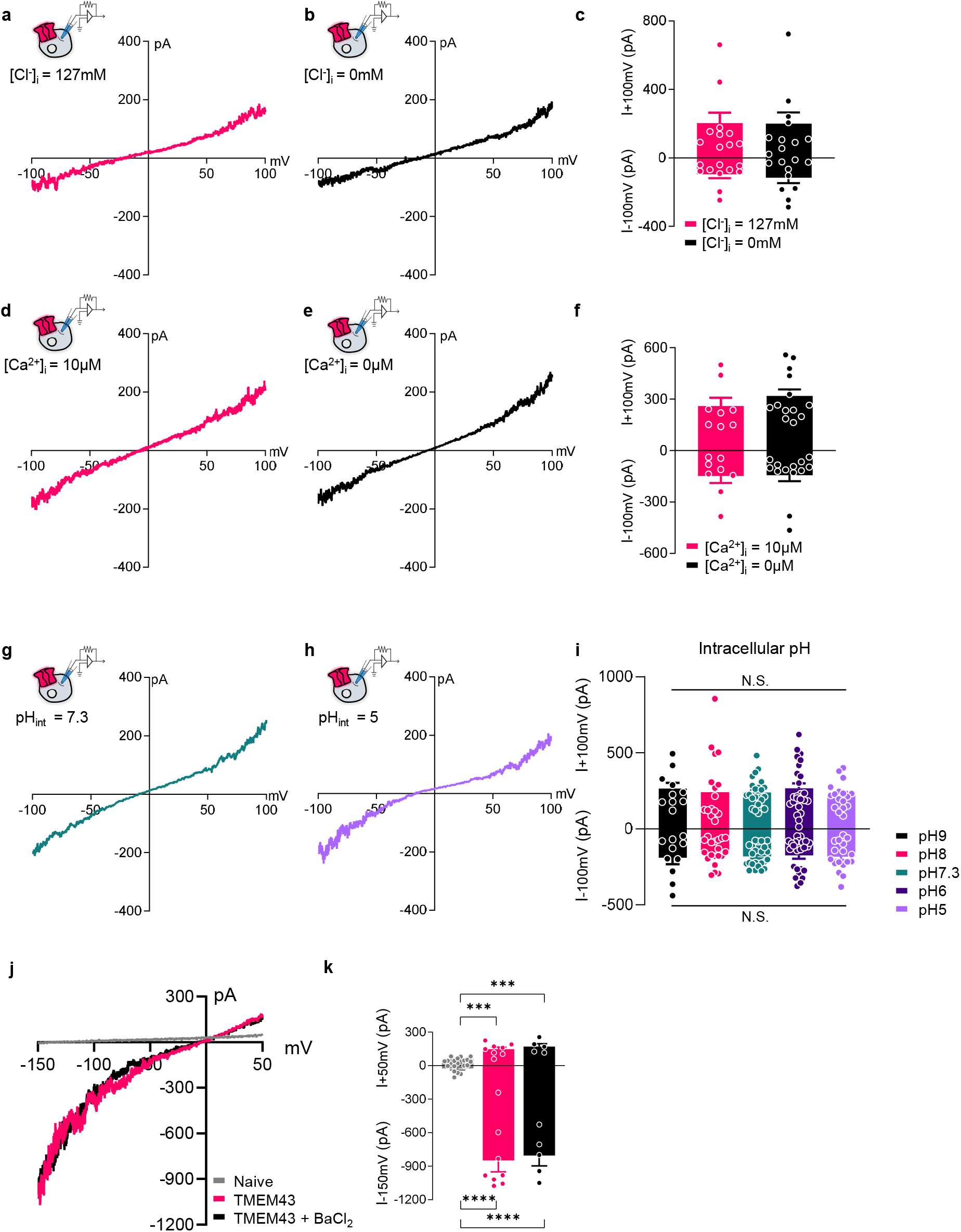
TMEM43-mediated current is independent of Cl^-^, Ca^2+^, internal pH changes, and Kir channels. **a,b,** Representative whole-cell I-V curves from HEK293T cells expressing TMEM43 with 127 mM [Cl^-^] (**a**) and 1 mM [Cl^-^] in the internal pipette solution (**b**). **c,** Summary bar graph of (**a,b**). **d,e,** Representative current traces from TMEM43-expressing cells with 10 μM [Ca^2+^] (**d**) and 0 μM [Ca^2+^] (**e**) in the patch pipettes. **f**, Summary bar graph of (**d,e**). **g,h,** Representative current-voltage relationship in TMEM43-expressing cells with internal pipette solution pH 7.3 (**g**) and pH 5 (**h**). **i,** Summary bar graph of TMEM43 currents measured with different pH in the internal solutions. **j**, Representative whole-cell I-V curves from naïve CHO-K1 cell and *TMEM43*-expressing cell before (pink) and after (black) BaCl_2_ treatment (100 μM shown). **k,** Summary bar graph of currents plotted at -150 mV (I_-150mV_) and +50 mV (I_+50mV_) for each condition in (**j**).

**Extended Data Fig. 2:**
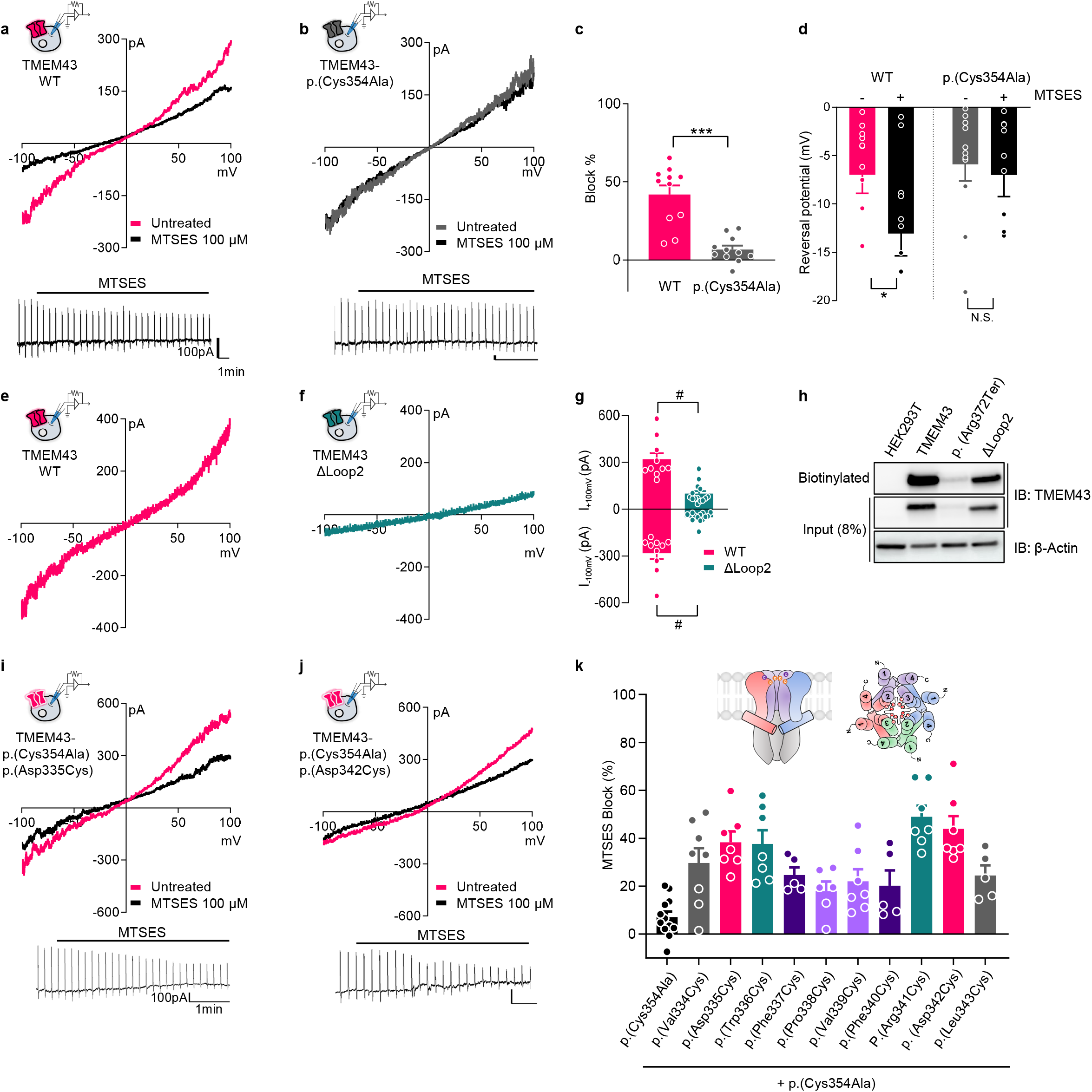
The putative channel pore domain of TMEM43 consists Loop2 domain. **a,b,** Representative I-V curves before (pink) and after (black) MTSES (100 μM) treatment in *TMEM43* WT-expressing (**a**) and *TMEM43-*p.(Cys354Ala)-expressing (**b**) cells. Raw traces are plotted in lower panels. **c,** Average current block percentage by MTSES treatment measured from (**a,b**). **d**, Average reversal potential measured from (**a,b**). **e,f**, Representative I-V curves of *TMEM43* WT (**e**) and Loop2 truncated (ΔLoop2) (**f**) -expressing cells. **g,** The average current amplitude of (**e,f**). **h**, Cell surface biotinylation assay in *TMEM43* WT and ΔLoop2 transfected HEK293T cells blotted with TMEM43 antibody. Naïve HEK293T cells and TMEM43-p.(Arg372Ter)-expressing cells were used as negative controls. **i,j**, Representative I-V curves before (pink) and after (black) 100 μM MTSES treatment of *TMEM43*-p.(Asp335Cys/Cys354Ala) transfected (**i**) and *TMEM43*-p.(Asp342Cys/Cys354Ala) transfected (**j**) cells. Raw traces are plotted in lower panels. (**k**) Average MTSES block percentage measured in cysteine substituted residues at Loop2 domain in p.(Cys354Ala) background. The proposed TMEM43 structure with putative channel pore is shown in the upper panel.

**Extended Data Fig. 3:**
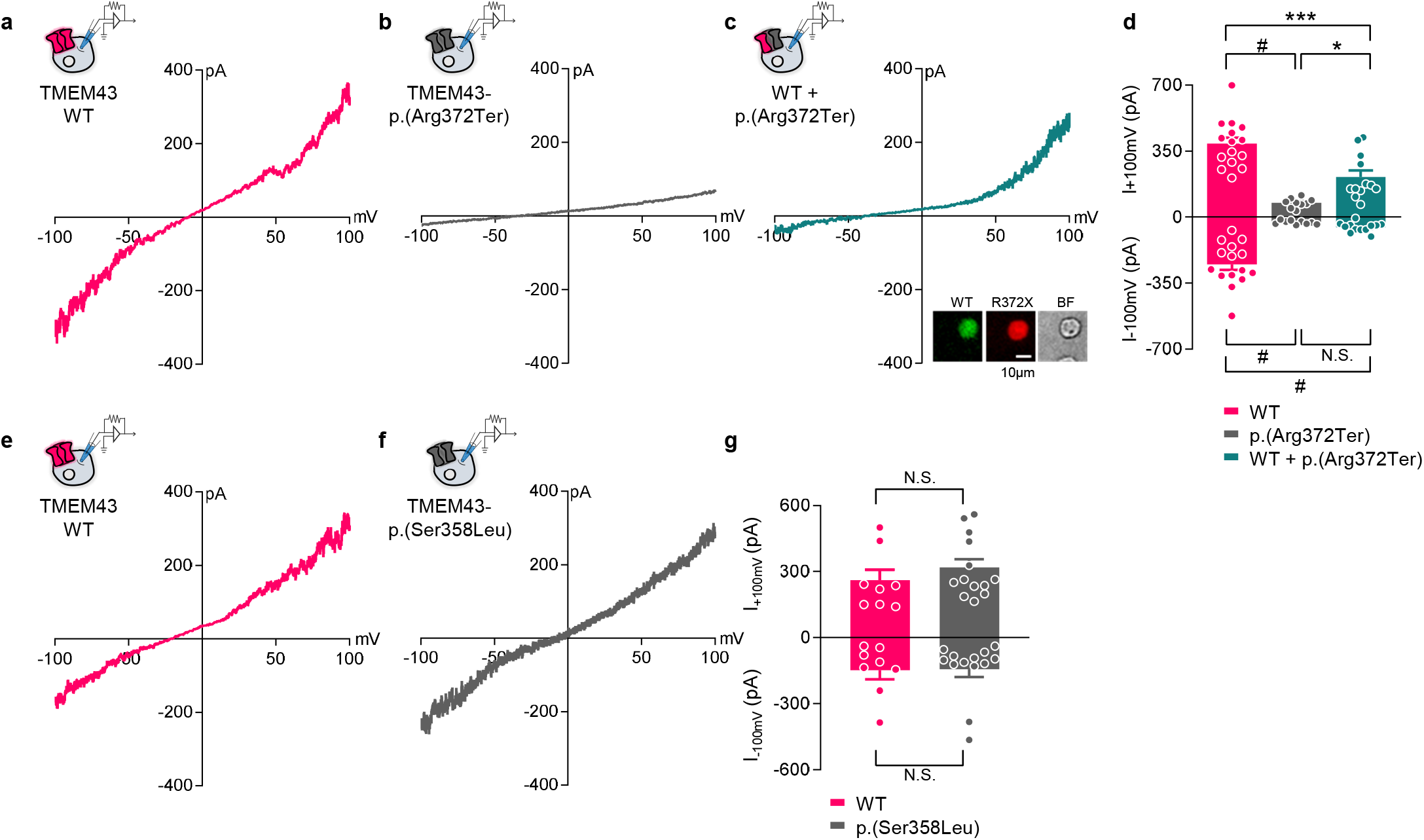
TMEM43-mediated currents with genetic variants. **a-c**, Representative whole-cell I-V curves from cells transfected with *TMEM43* WT (**a**), *TMEM43*-p.(Arg372Ter) (**b**), or *TMEM43* WT and *TMEM43*-p.(Arg372Ter) (1:1) (**c**). Insets: fluorescent images of a CHO-K1 cell expressing both TMEM43 WT (GFP) and TMEM43-p.(Arg372Ter) (tdTomato). **d,** Summary bar graph of currents from (**a-c**). **e-f,** Representative whole-cell I-V curves from cells transfected with *TMEM43* WT (**e**) and *TMEM43*-p.(Ser358Leu) (**f**). **g,** Averaged current amplitudes from (**e,f**).

**Extended Data Fig. 4:**
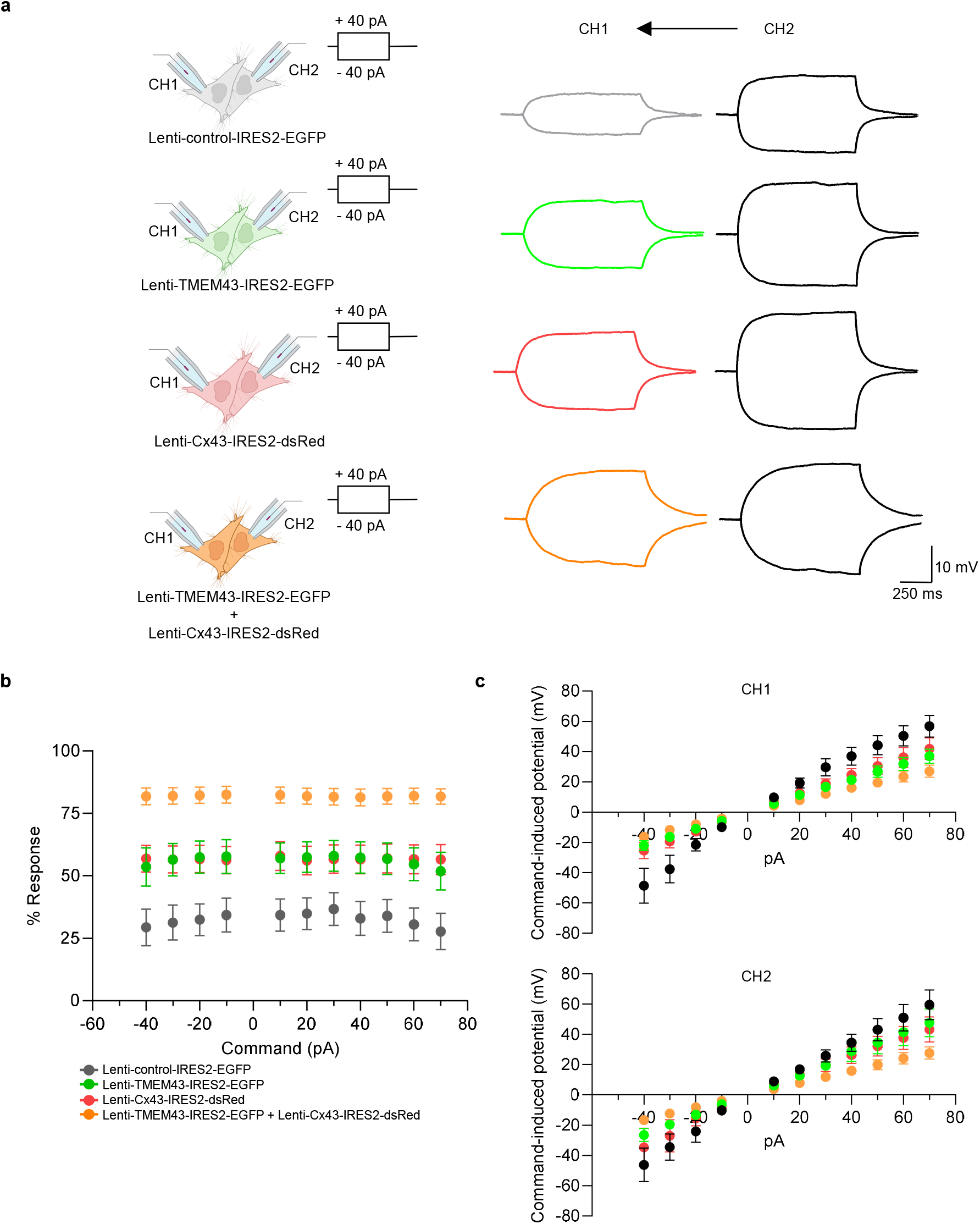
TMEM43 mediated electrical couplings in HeLa cells. **a,** Representative traces showing electrical couplings from sender cells (black traces) to receiver cells (gray and color traces) in different lentivirus-infected HeLa cells. **b**, Averaged % responses of electrical coupling on receiver cells (CH1) at various commands from sender cells (CH2). **c**, Averaged potential of receiver cells induced by different commands from sender cells at different conditions.

**Extended Data Fig. 5:**
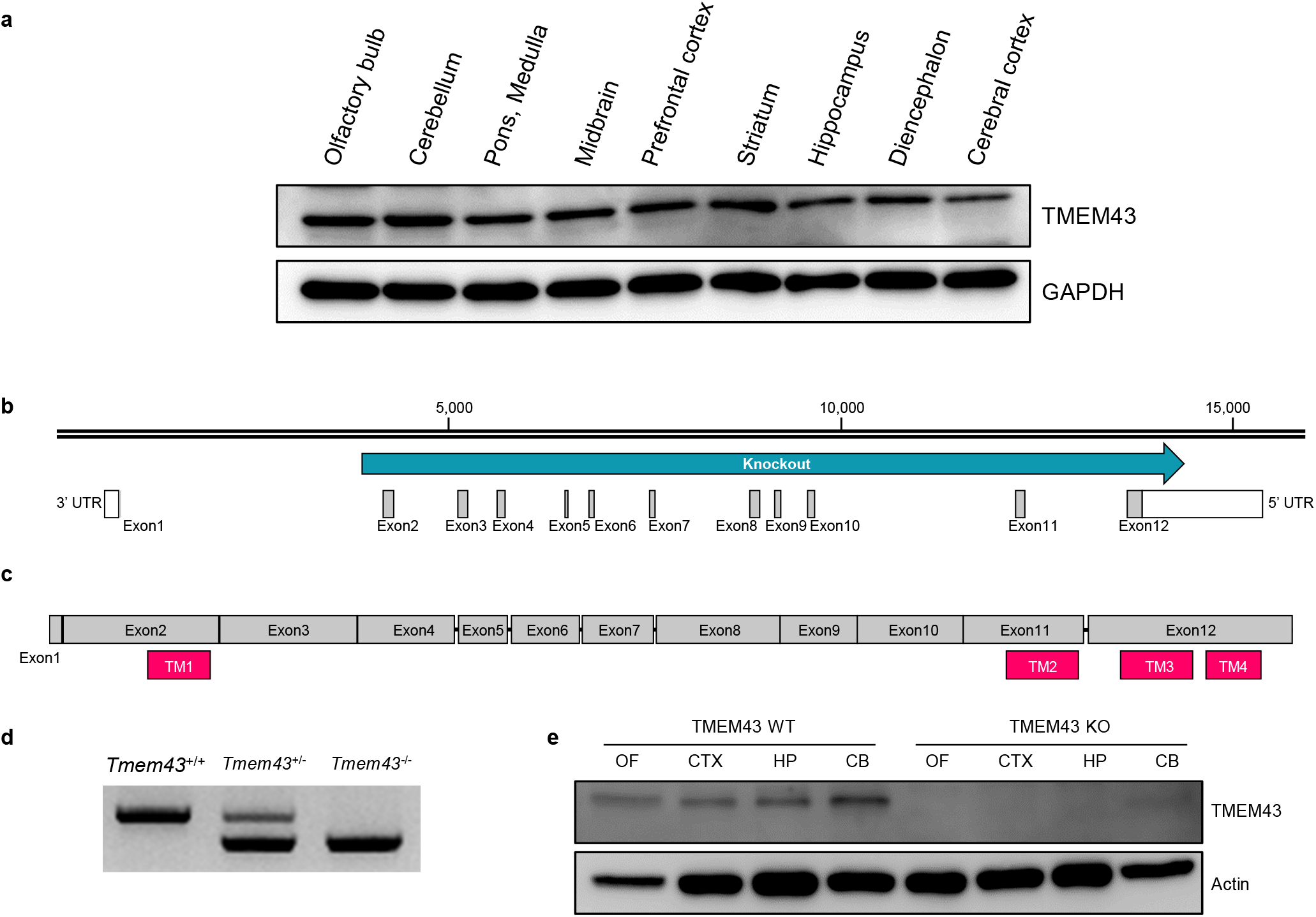
TMEM43 expression in the brain and validation of TMEM43 KO mice. **a,** Western blot assay depicting TMEM43 expression in nine different brain regions. **b**, Illustration of TMEM43 exon map on mouse genome. Knockout region is indicated in blue. **c**, Illustration of TMEM43 exon map after splicing. TM domains are indicated in pink. **d**, Genotyping result of *Tmem43^+/+^*, *Tmem43^+/-^*, and *Tmem43^-/-^*. **e**, Western blot assay from brain tissue lysates of TMEM43 WT (left) and KO (right). OF: olfactory bulb, CTX: cortex, HP: hippocampus, CB: cerebellum.

**Extended Data Fig. 6:**
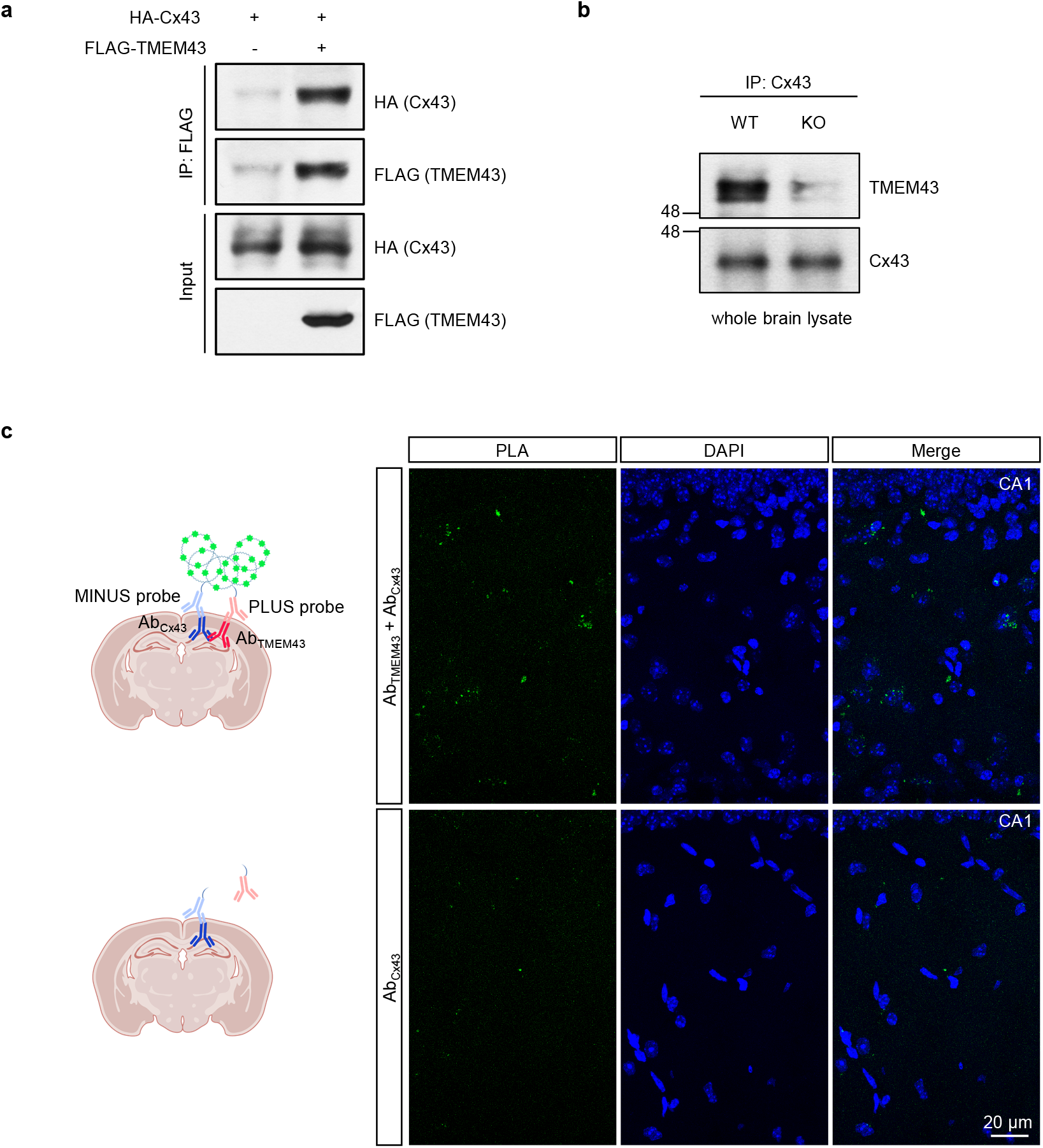
TMEM43 interacts with Cx43 gap junction channels. **a,b**, Co-IP data showing that TMEM43 is immuno-pulled down with Cx43 in *in vitro* (**a**) and in brain lysates (**b**). **c**, PLA results with anti-TMEM43 and anti-Cx43 (Top) done in hippocampal brain slices. PLA signal was amplified as a red fluorescent, indicative of close proximity of TMEM43 and Cx43. Red signal was pseudo colored as green for better data display. Lower panel is a negative control data without TMEM43 antibody.

**Extended Data Fig. 7:**
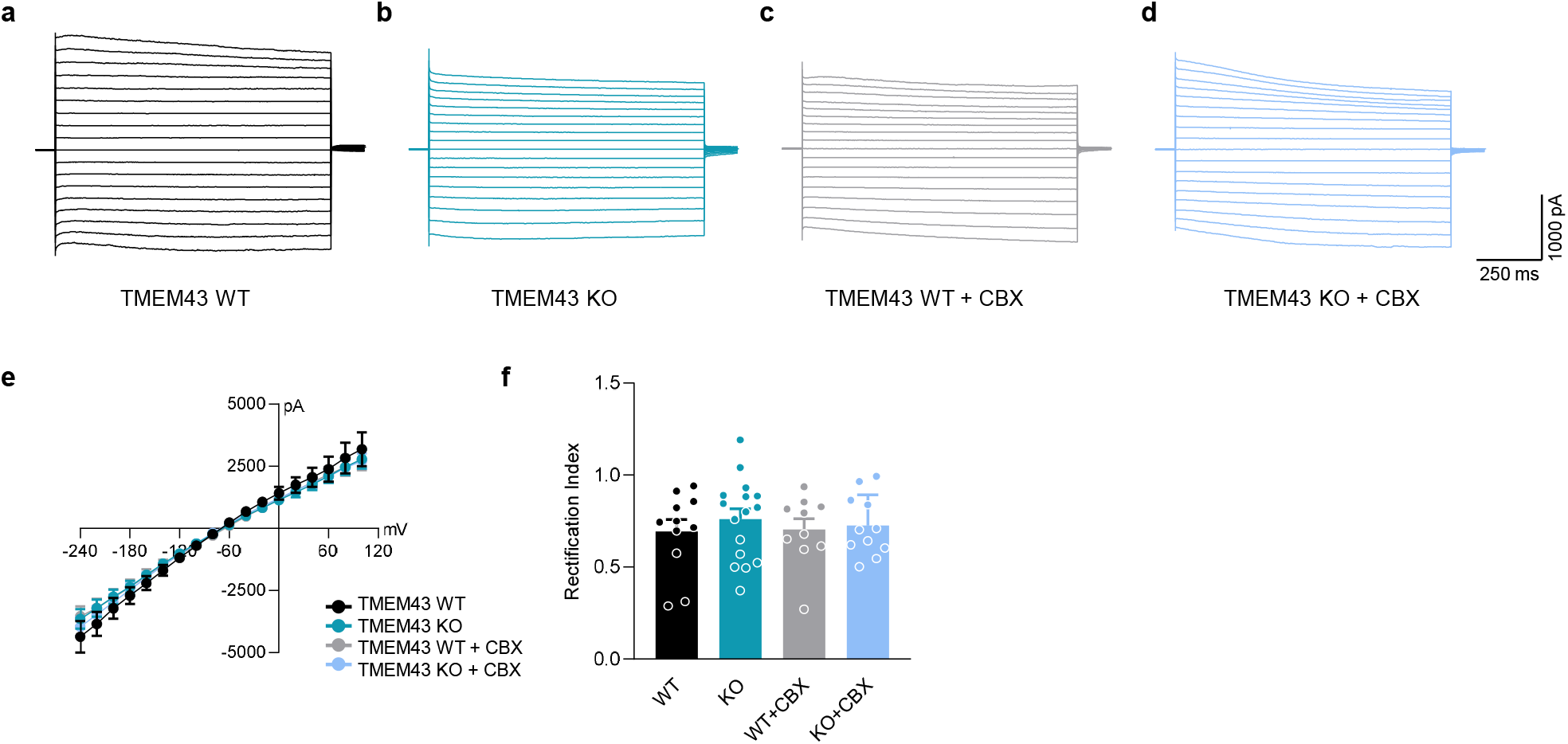
Astrocytic passive conductance current is not disturbed in TMEM43 KO mice. **a-d**, Representative traces of astrocytic passive conductance current measured in hippocampal striatum radiatum of TMEM43 WT (**a**), TMEM43 KO (**b**), TMEM43 WT with CBX treatment (**c**), and TMEM43 KO with CBX treatment (**d**). **e**, Averaged current-voltage relationship of hippocampal striatum radiatum astrocytes at different conditions. **F**, Rectification index measured from each group.

**Extended Data Fig. 8:**
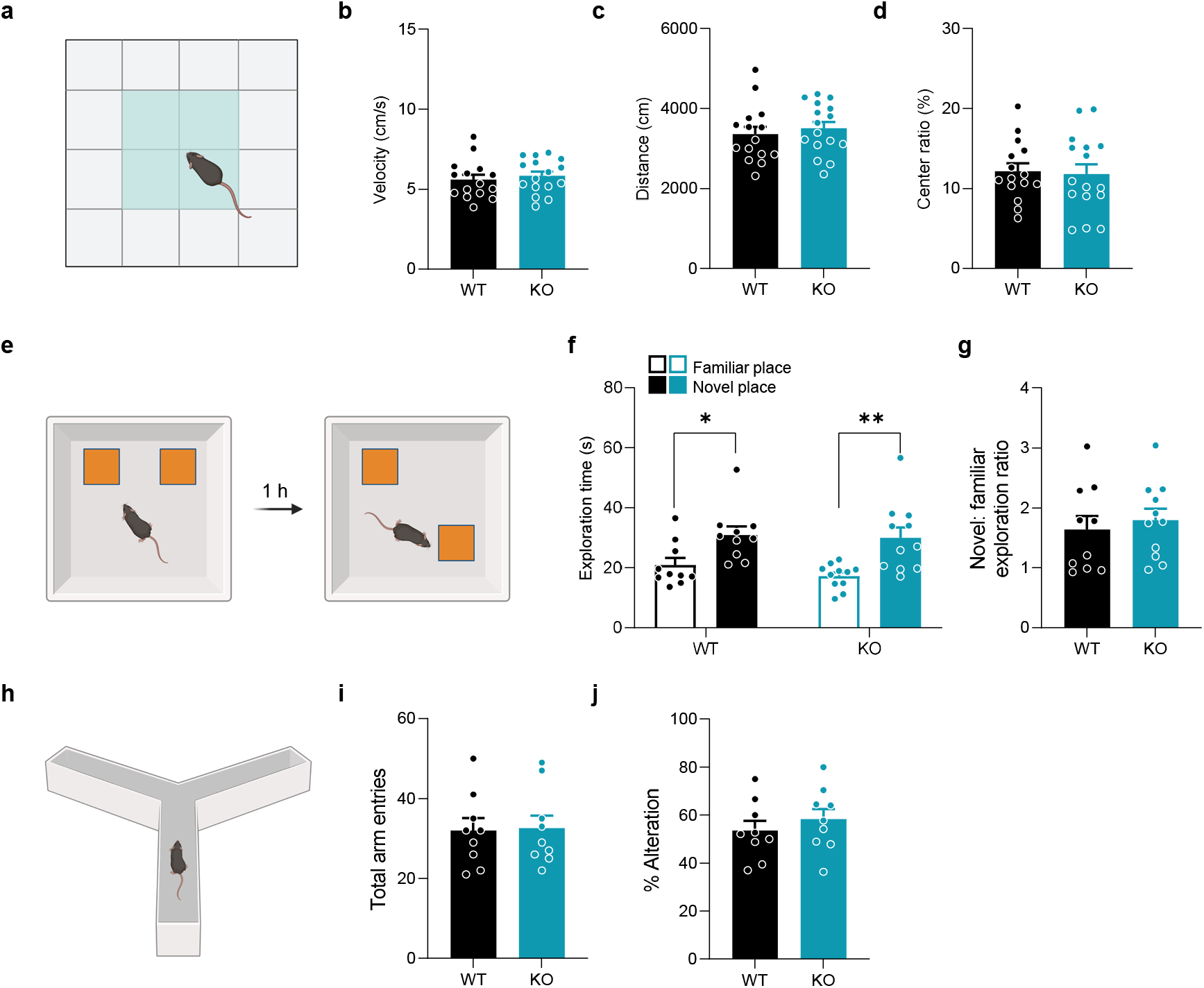
General behavior assays of TMEM43 KO mouse line. **a,** Open field test. Shaded area in the middle is considered as center. **b-d**, Averaged velocity (**b**), distance moved (**d**), and center ratio (**d**) from TMEM43 WT (black) and KO (blue). **e**, Experimental scheme of novel place recognition test. Subject mouse was allowed to explore two identical objects for 10 min. After 1 hour, one of the two identical objects was placed at a novel place and exploration time was measured. **f**, Averaged exploration time compared between familiar (blank bar) and novel (filled bar) place from TMEM43 WT (black) and KO (blue). **g**, Novel: familiar exploration ratio form (**f**). **h**, Y-maze with three open arms. **i**,**j**, Averaged total arm entries (**i**) and % alteration (**j**) from TMEM43 WT (black) and KO (blue).

## Notes

### Competing Interest Statement

The authors have declared no competing interest.

### Summary of Updates

Functional studies have been added.

